# Thiol drugs decrease SARS-CoV-2 lung injury *in vivo* and disrupt SARS-CoV-2 spike complex binding to ACE2 *in vitro*

**DOI:** 10.1101/2020.12.08.415505

**Authors:** Kritika Khanna, Wilfred Raymond, Jing Jin, Annabelle R. Charbit, Irina Gitlin, Monica Tang, Adam D. Werts, Edward G. Barrett, Jason M. Cox, Sharla M. Birch, Rachel Martinelli, Hannah S. Sperber, Sergej Franz, Satish Pillai, Anne Marie Healy, Thomas Duff, Stefan Oscarson, Markus Hoffmann, Stefan Pöhlmann, Graham Simmons, John V. Fahy

## Abstract

Neutrophil-induced oxidative stress is a mechanism of lung injury in COVID-19, and drugs with a functional thiol group (“thiol drugs”), especially cysteamine, have anti-oxidant and anti-inflammatory properties that could limit this injury. Thiol drugs may also alter the redox status of the cysteine-rich SARS-CoV-2 spike glycoprotein (SARS-2-S) and thereby disrupt ACE2 binding. Using ACE2 binding assay, reporter virus pseudotyped with SARS-CoV-2 spikes (ancestral and variants) and authentic SARS-CoV-2 (Wuhan-1), we find that multiple thiol drugs inhibit SARS-2-S binding to ACE2 and virus entry into cells. Pseudoviruses carrying variant spikes were less efficiently inhibited as compared to pseudotypes bearing an ancestral spike, but the most potent drugs still inhibited the Delta variant in the low millimolar range. IC50 values followed the order of their cystine cleavage rates and lower thiol pKa values. In hamsters infected with SARS-CoV-2, intraperitoneal (IP) cysteamine decreased neutrophilic inflammation and alveolar hemorrhage in the lungs but did not decrease viral infection, most likely because IP delivery could not achieve millimolar concentrations in the airways. These data show that thiol drugs inhibit SARS-CoV-2 infection *in vitro* and reduce SARS-CoV-2-related lung injury *in vivo* and provide strong rationale for trials of systemically delivered thiol drugs as COVID-19 treatments. We propose that antiviral effects of thiol drugs *in vivo* will require delivery directly to the airways to ensure millimolar drug concentrations and that thiol drugs with lower thiol pKa values are most likely to be effective.

**One Sentence Summary:** The effect of cysteamine to decrease SARS-CoV-2 pneumonia *in vivo* and of multiple thiol drugs to inhibit SARS-CoV-2 infection *in vitro* provides rationale for clinical trials of thiol drugs in COVID-19.

## INTRODUCTION

SARS-CoV-2 is a novel coronavirus that causes COVID-19, a multidimensional disease characterized predominantly by pneumonia that can progress to respiratory failure and death (*1, 2*). Patients with severe COVID-19-related pneumonia have high neutrophil to lymphocyte ratios and high levels of reactive oxygen species (ROS) which cause lung injury (*3*). Multiple anti-inflammatory treatments, including corticosteroids and cytokine inhibitors have been used to treat SARS-CoV-2-related lung injury (*4–7*), and corticosteroids have been especially effective, most likely because of their broad anti-inflammatory effects (*8, 9*). Drugs that have at least one functional thiol group (“thiol drugs”, Table 1) also have broad antioxidant and anti-inflammatory effects. Specifically, thiols drugs scavenge reactive oxygen species (ROS) and interrupt ROS-mediated inflammatory cascades, including pathways for NFkB activation and cytokine and chemokine production (*10–15*), and can inhibit the activity of myeloperoxidase, a mediator of inflammation and oxidative stress (*16, 17*). Cysteamine is a thiol drug with an expanded range of anti-inflammatory activities related to inhibition of transglutaminase 2 and somatostatin (*18, 19*). Cysteamine decreases aeroallergen-induced lung inflammation in mice (*20, 21*) which raises the possibility that it could decrease SARS-CoV-2-induced lung inflammation as well. Golden Syrian hamsters (Mesocricetus auratus) infected with SARS-CoV-2 develop diffuse alveolar destruction, airway and alveolar infiltration with inflammatory cells (macrophages, lymphocytes, and neutrophils) and alveolar hemorrhage (*22–26*). The hamster model is therefore a valuable model to test if drugs have therapeutic potential against SARS-CoV-2 pneumonia.

**Table 1:**
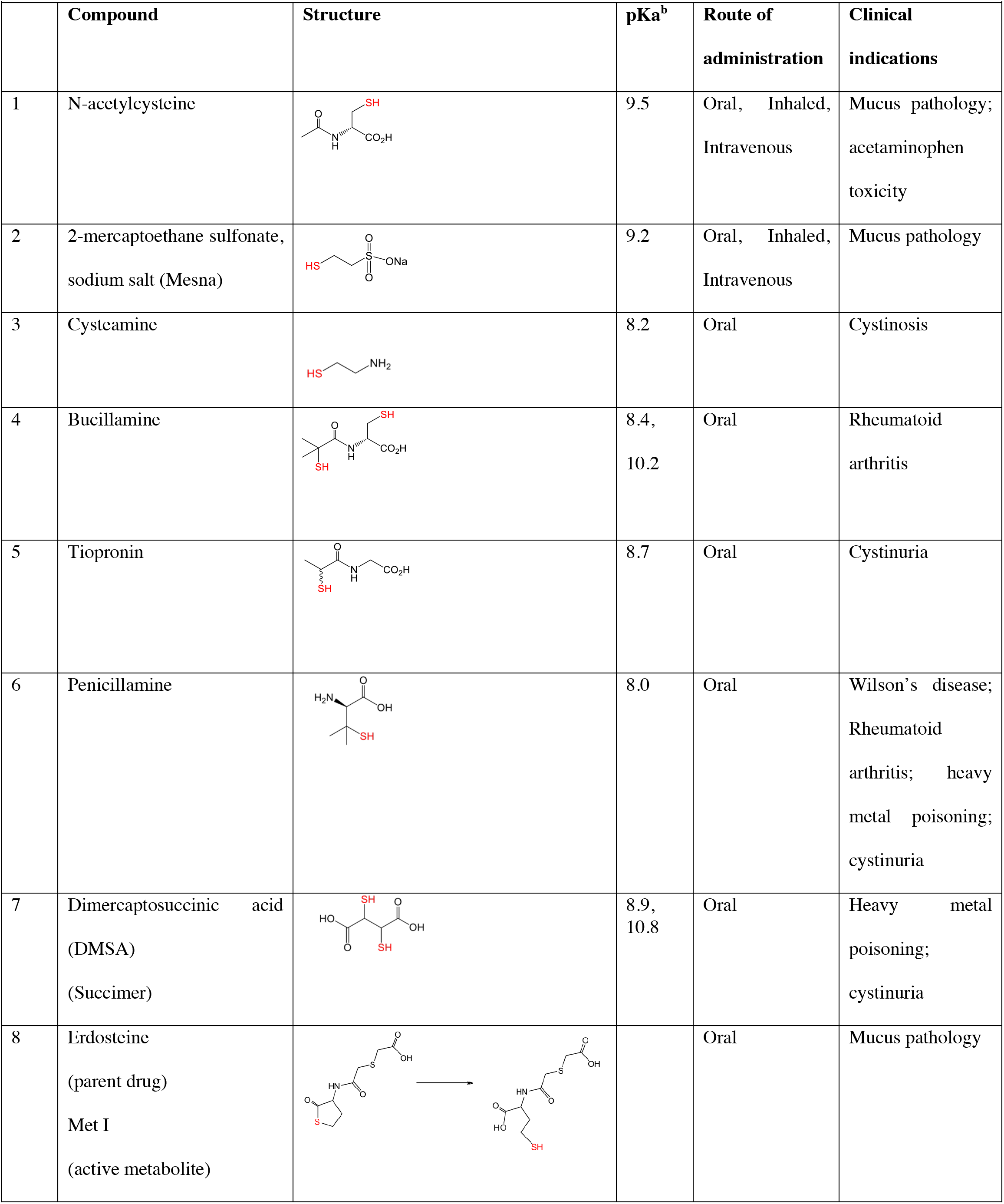
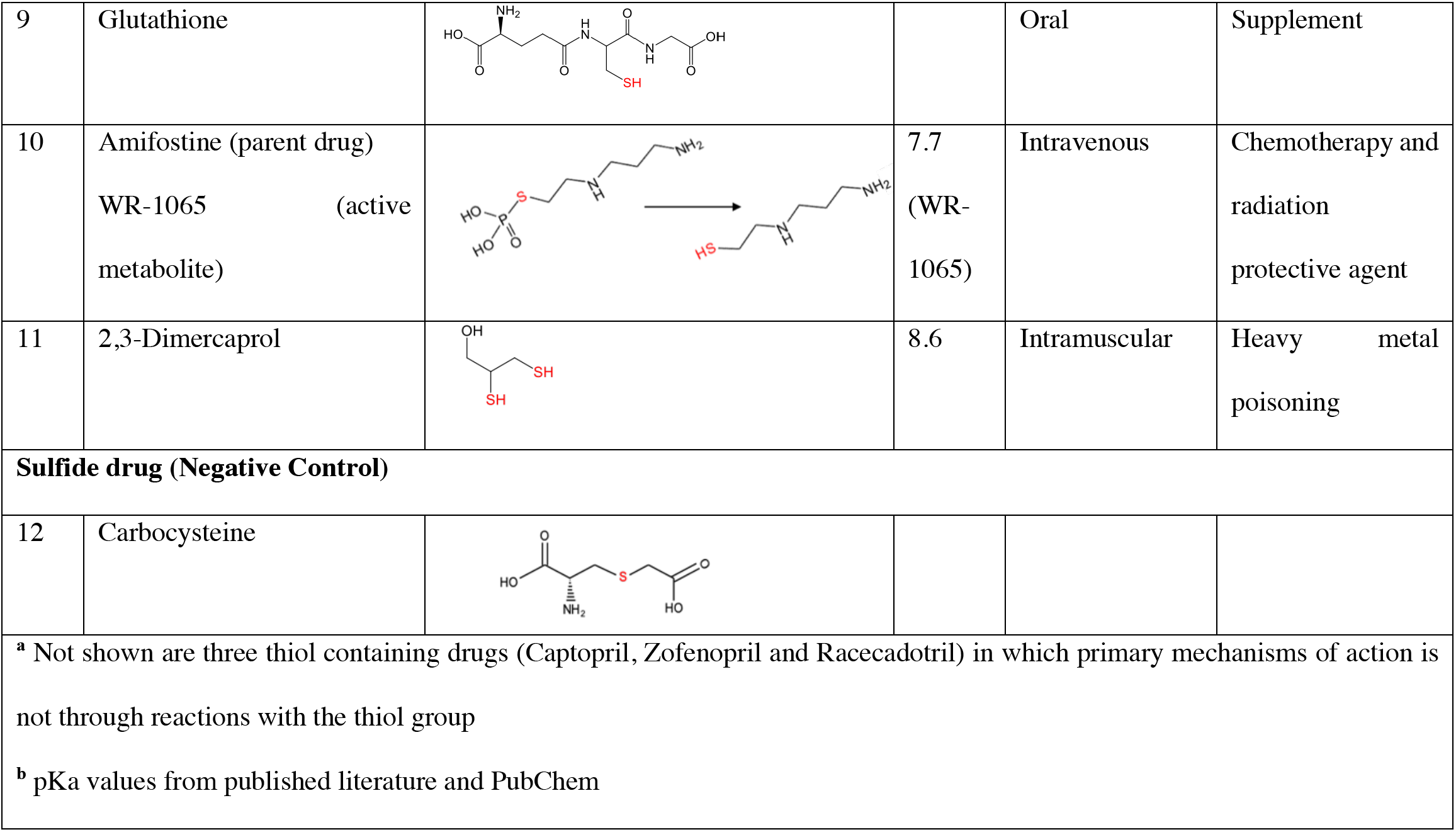
Comprehensive list of currently approved thiol-based drugs or drugs that generate a thiol-containing metabolite^a^.

|  |  |  |  |  |  |
| --- | --- | --- | --- | --- | --- |
|  | Compound | Structure | pK<sub>a</sub><sup>b</sup> | Route of administration | Clinical indications |
| 1 | N-acetylcysteine |  | 9.5 | Oral, Inhaled, Intravenous | Mucus pathology; acetaminophen toxicity |
| 2 | 2-mercaptoethane sulfonate, sodium salt (Mesna) |  | 9.2 | Oral, Inhaled, Intravenous | Mucus pathology |
| 3 | Cysteamine |  | 8.2 | Oral | Cystinosis |
| 4 | Bucillamine |  | 8.4, 10.2 | Oral | Rheumatoid arthritis |
| 5 | Tiopronin |  | 8.7 | Oral | Cystinuria |
| 6 | Penicillamine |  | 8.0 | Oral | Wilson's disease; Rheumatoid arthritis; heavy metal poisoning; cystinuria |
| 7 | Dimercaptosuccinic acid (DMSA) (Succimer) |  | 8.9, 10.8 | Oral | Heavy metal poisoning; cystinuria |
| 8 | Erdosteine (parent drug) Met I (active metabolite) |  |  | Oral | Mucus pathology |
| 9 | Glutathione |  |  | Oral | Supplement |
| 10 | Amifostine (parent drug)<br>WR-1065 (active metabolite) |  | 7.7<br>(WR-1065) | Intravenous | Chemotherapy and radiation protective agent |
| 11 | 2,3-Dimercaprol |  | 8.6 | Intramuscular | Heavy metal poisoning |
| **Sulfide drug (Negative Control)** |  |  |  |  |  |
| 12 | Carbocysteine |  |  |  |  |
| <sup>a</sup> Not shown are three thiol containing drugs (Captopril, Zofenopril and Racecadotril) in which primary mechanisms of action is not through reactions with the thiol group<br><sup>b</sup> pKa values from published literature and PubChem |  |  |  |  |  |

The envelope glycoproteins of SARS-CoV-2 form trimeric spikes (SARS-2-S) that bind the cell surface receptor, angiotensin converting enzyme 2 (ACE2) for virus entry (*27*). Coronavirus spike glycoproteins, together with the envelope glycoproteins of myxo-and paramyxoviruses, retroviruses and filoviruses belong to class I viral fusion proteins that exhibit similar structural features and mechanistic strategies to fuse viral and cellular membranes, which is critical for infectious viral entry (*28*). The capacity of class I viral fusion proteins to mediate membrane fusion often depends on a precise thiol/disulfide balance in the viral fusion proteins (*29–32*). Natural and specific thiol/disulfide rearrangements in the fusion proteins can trigger conformational changes that promote virus entry (*33–35*), but removal of disulfide bridges by chemical reduction or by mutation of cysteines can also disrupt virus binding to receptors and therefore prevent infection. For example, chemical reduction of the S1 domain of SARS-CoV decreases its binding to ACE2 and inhibits infection of Vero E6 cells by SARS-CoV pseudoviruses, and replacing cystine forming cysteines with alanines in the receptor binding domain (RBD) of SARS-CoV spike (hereafter SARS-1-S) prevents its binding to ACE2 (*32*). In addition, molecular dynamics simulations reveal that the binding affinity of SARS-2-S RBD for ACE2 is significantly impaired when all of the disulfide bonds of both ACE2 and SARS-2-S are reduced to thiol groups (*36*). Thus, there is consistent literature that manipulation of the redox status of the cysteine-rich glycoproteins on the virus surface can influence or impair viral infectivity. This led us to consider that thiol drugs may cleave cystines in the RBD of SARS-2-S to disrupt its binding to ACE2. The reactive or cystine cleaving form of a thiol is its deprotonated (thiolate) form (*37*), and the pKa or acid dissociation constant of a thiol group determines the fractions of the protonated (-SH) form and the deprotonated (-S^-^) forms at the pH of the working solution. Alkyl thiols are weakly acidic with pKa >9, with low fraction of thiolate present at physiologic pH of 7.4. Thiol compounds with a lower pKa (i.e. approaching 7.4) will have more thiolate species at physiologic conditions and may be more potent in inhibition of SARS-CoV-2 infection.

In this study, we screened the efficacy of thiol drugs in SARS-2-S mediated receptor binding and virus infection assays, and we explored the role of thiol pKa as a physicochemical property of thiol drug potency in these assays. In *in vivo* studies in hamsters, we tested if cysteamine decreases the severity of SARS-CoV-2-induced lung injury.

## RESULTS

### Cystine bridges in the SARS-2-S1 domain

It is not known if thiol drugs can disrupt cystines in SARS-2-S to inhibit binding to ACE2 or virus entry into cells. To begin to address these questions, we first used published data (*38, 39*) to build a cystine bridge map of the S1 domain of SARS-2-S, and we compared the amino acid alignment of the RBDs in SARS-2-S and SARS-1-S. We noted 10 cystine bridges in the SARS-2-S1 domain (Fig. 1A) and 4 conserved cystines between SARS-1-S and SARS-2-S RBD (Fig. 1B). The conserved Cys467-Cys474 in SARS-1-S and Cys480-Cys488 in SARS-2-S constrain the ACE2 binding domains, and previous studies with SARS-1-S RBD have shown that mutagenesis of either conserved cysteine leads to loss of ACE-2 binding(*32*). To further explore if Cys480-Cys488 in SARS-2-S might be vulnerable to chemical cleavage, we used protein modeling software to render the SARS-2-S RBD based on PDB entry 6M0J (Fig. 1C). This rendering shows that Cys480-Cys488 is very near the RBD surface (Fig. 1C) and could be accessible to cleavage by thiol-based drugs. Apart from Cys480-Cys488 cystine, cleavage of the other six cystines in the RBD could also allosterically modify the binding interface in ways that decrease its binding to ACE2. Studies in SARS-CoV have shown that cysteine residues flanking the S2 domain - including Cys822 and Cys833 - are important for membrane fusion of SARS-CoV (*40*). Amino acid alignment of the spike protein of SARS-CoV and SARS-CoV-2 shows that these cysteine residues are conserved in the spike protein of SARS-CoV-2 (Cys 840 and Cys 851) (data not shown), which raises the possibility that thiol-based drugs could inhibit membrane fusion in addition to inhibitory effects on receptor binding.

**Fig. 1.**
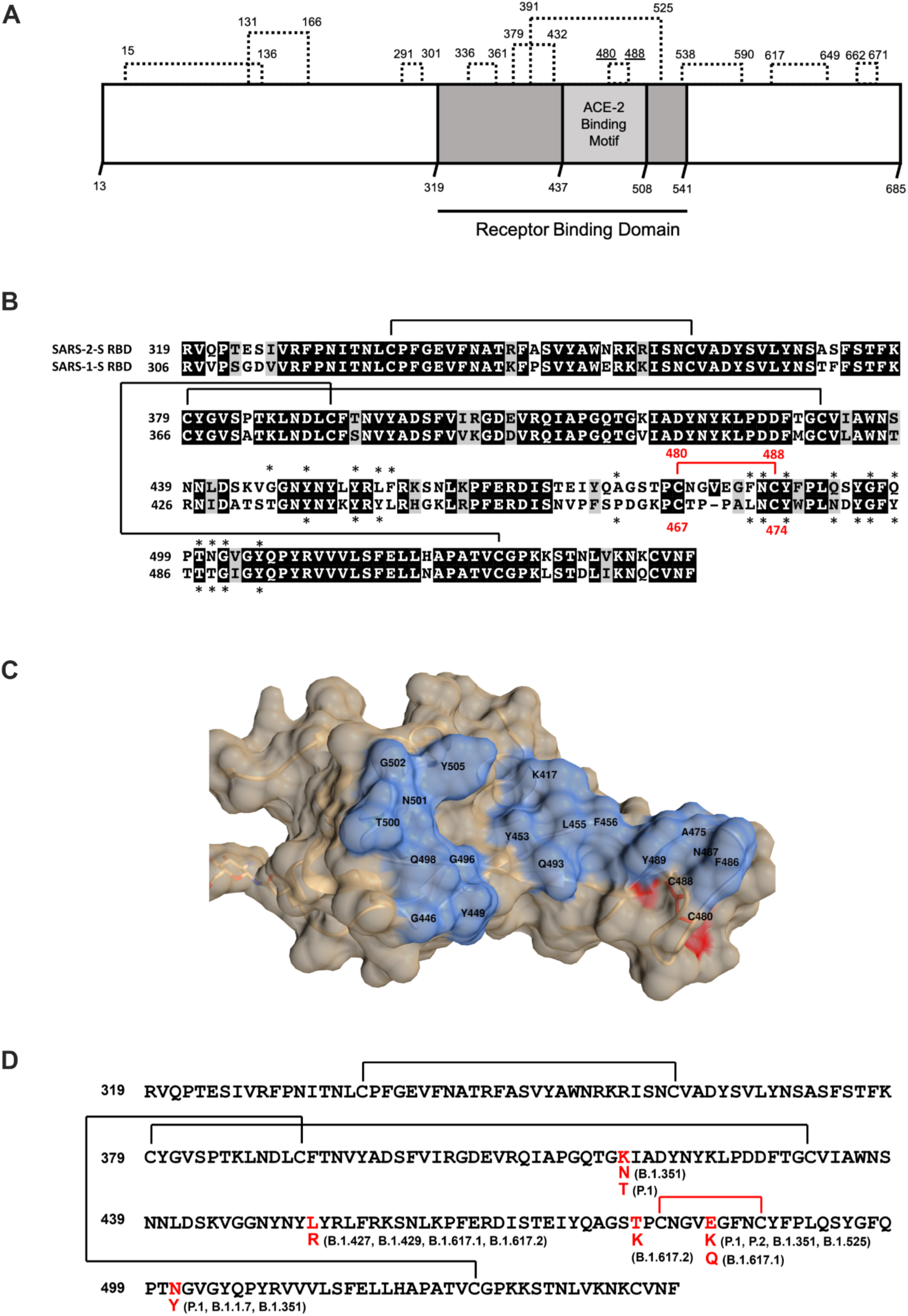
Cystine mapping and conservation of cystines in RBD of SARS-1-S and SARS-2-S. (**A**) Cystine map for SARS-2-S domain S1 (UniProt: P0DTC2). Ten cystine linkages are denoted by dashed lines with amino acid residue number above. The dark gray region is the RBD, and the lighter gray box highlights the ACE2 binding motif, a cluster of amino acids that make contact with ACE2. (**B**) Amino acid alignment of SARS-2-S RBD domain (aa 319-541, PDB: 6M0J) and SARS-1-S RBD domain (aa 306-517, PDB: 3SCI). Residues that are shared are highlighted by black boxes and residues that represent a similar amino acid class replacement are bound by gray boxes. The solid lines link cystine-forming cysteines. The solid red line and red numbers highlight the conserved critical cystine bridge in the RBDs for both viruses. Asterisks denote amino acids that are within 4 angstroms of ACE2 in their respective solved structures. (**C**) A surface rendering of SARS-2-RBD (PDB: 6M0J) oriented with the ACE2 binding region (blue) facing forward. (**D**) Amino acid sequence of SARS-2-S RBD highlighting the RBD mutations identified in the circulating SARS-CoV-2 variants. Amino acids are noted with single letter code and sequence number. The conserved RBD cystine (C480-C488) is highlighted in red.

Like other RNA viruses, SARS-CoV-2 undergoes random genome mutations that are subject to natural selection. Consequently, SARS-CoV-2 variants have emerged with amino acid substitutions and deletions or combinations of substitutions in the spike glycoprotein. Some variants are classified as variants of interest (VOI) or variants of concern (VOC) due to their increased transmissibility and/or evasion from natural or therapeutic antibodies (*41*). In particular, the highly infectious Delta variant has spread rapidly and is now globally dominant (*42, 43*). We annotated point mutations and displayed the intact cystine bridges in the RBD of SARS-CoV-2 VOI and VOC in Fig. 1D.

### Thiol drugs cleave cystines in the RBD of SARS-2-S to disrupt binding to ACE2

To test if thiol drugs cleave cystines in the RBD of SARS-2-S to disrupt its binding to ACE2, we exposed the RBD of the ancestral SARS-CoV-2 Wuhan-1 isolate (RBD^Wuhan-1^) to 8 thiol drugs and quantified ACE2 binding affinity in a plate-based binding assay. Carbocysteine and amifostine were included as negative controls because carbocysteine is a sulfur-containing drug lacking a free thiol warhead and amifostine is a phosphorothioate prodrug whose dephosphorylated metabolite (WR-1065) is the active form of the drug (Table 1). A commercially available ACE2-SARS-2-S RBD binding assay was optimized by covalently coupling RBD^Wuhan-1^ to plates functionalized with primary amine-reactive maleic anhydride, and ACE2 binding was quantified after RBD^Wuhan-1^ exposure to drugs for 60 minutes (Fig. 2A). We found that all of the thiol drugs inhibited binding of RBD^Wuhan-1^ to ACE2 in a dose dependent manner. Penicillamine and succimer had relatively weak inhibitory effects (fig. S1), but 2-mercaptoethane sulfonate sodium salt (Mesna), bucillamine, cysteamine and WR-1065 had much stronger effects (Fig. 2B, C). To determine if this binding inhibition is retained after drug removal, ACE2 binding to RBD^Wuhan-1^ was measured two hours after exposure to the most potent thiol drugs. In this way, we showed that binding inhibition was retained for two hours (Fig. 2D).

**Fig. 2.**
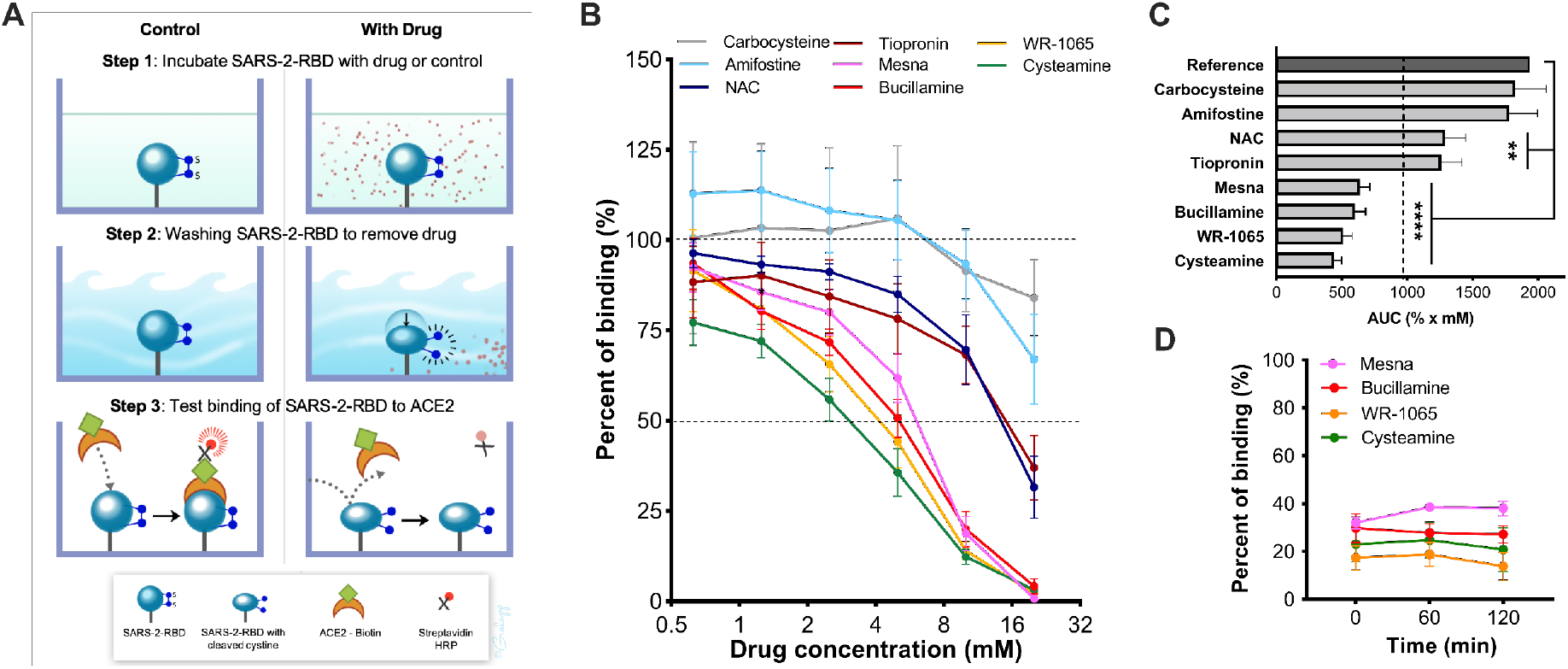
Binding of SARS-CoV-2 RBD to ACE2 is inhibited by thiol-based drugs. (**A**) Schematic representation of the SARS-CoV-2 RBD to ACE2 binding assay. (**B**) Percent of binding of RBD^Wuhan-1^ to ACE2 in the presence of the drugs (n = 4-6). (**C**) Area under the curve (AUC) analysis of (B). (**D**) Binding of RBD^Wuhan-1^ to ACE2 at one and two hours post drug exposure and washout (n = 4 - 5). Reference AUC was calculated from RBD to ACE2 binding with no drug control; dashed line represents 50% of reference AUC. Statistical significance for (C) was analyzed by one-way ANOVA followed by Dunnett’s post-hoc analysis. Significance indicates differences from reference AUC. **p ≤ 0.01, ****, p ≤ 0.0001.

### Thiol drugs inhibit infection with SARS-2-S bearing pseudotypes and authentic SARS-CoV-2

We employed vesicular stomatitis virus (VSV)-derived pseudovirus (PV) particles carrying SARS-2-S on their surface and harboring a reporter genome with a deleted glycoprotein gene (rVSV-ΔG) and an insertion of the firefly luciferase gene (*44*). To test if thiol drugs inhibit SARS-CoV-2 pseudovirus (SARS-2-PV) entry efficiency, we first treated SARS-2-PV particles bearing spike from ancestral strain, Wuhan-1, with thiol-based drugs and quantified their infectivity using 293T-ACE2-TMPRSS2 cells, human embryonic kidney cell line (HEK293T) stably expressing human ACE2 and transmembrane protease, serine 2 (TMPRSS2, a priming serine protease for SARS-CoV-2) (*27*). As illustrated in fig. S2, the experimental protocol measured SARS-2-PV entry into 293T-ACE2-TMPRSS2 cells with either the virus or the target cells being pre-treated with thiol-based drugs for 2 hours prior to transduction. None of the drugs significantly affected cell viability, and pretreatment of SARS-2-PV with carbocysteine and amifostine controls did not inhibit virus entry into the cells (Fig. 3A, B). In contrast, pretreatment of SARS-2-PV with thiol drugs significantly inhibited virus entry in a dose dependent manner (Fig. 3C-H). Cysteamine and WR-1065 were much more effective than NAC and tiopronin (Fig. 3I). Thiol drugs had only small and inconsistent effects on PV infectivity when the 293T-ACE2-TMPRSS2 cells were pretreated with thiol drugs and then infected with untreated SARS-2-PV (fig. S3A-H). To confirm that these data with PV particles extend to authentic SARS-CoV-2, we initially carried out dose ranging experiments to preliminarily test the effects of the most potent thiol drugs (Mesna, bucillamine, WR-1065 and cysteamine) on SARS-CoV-2 (Wuhan-1) infection in Vero E6 cells. We found that cysteamine and WR-1065 were very effective at decreasing virus-induced cytopathic effects (fig. S4A-E). In a subsequent optimized experiment, we tested the effects of cysteamine on SARS-CoV-2 infection of Vero E6 cells stably expressing TMPRSS2 (Vero E6-TMPRSS2). We found that cysteamine significantly decreased virus-induced cytopathic effects whereas carbocysteine was not effective (Fig. 3J, K). For both preliminary and optimized experiments, inhibition of infection was minimal when the respective cells were first pretreated with the thiol drugs and then infected with untreated SARS-CoV-2 (fig. S4F, G).

**Fig. 3.**
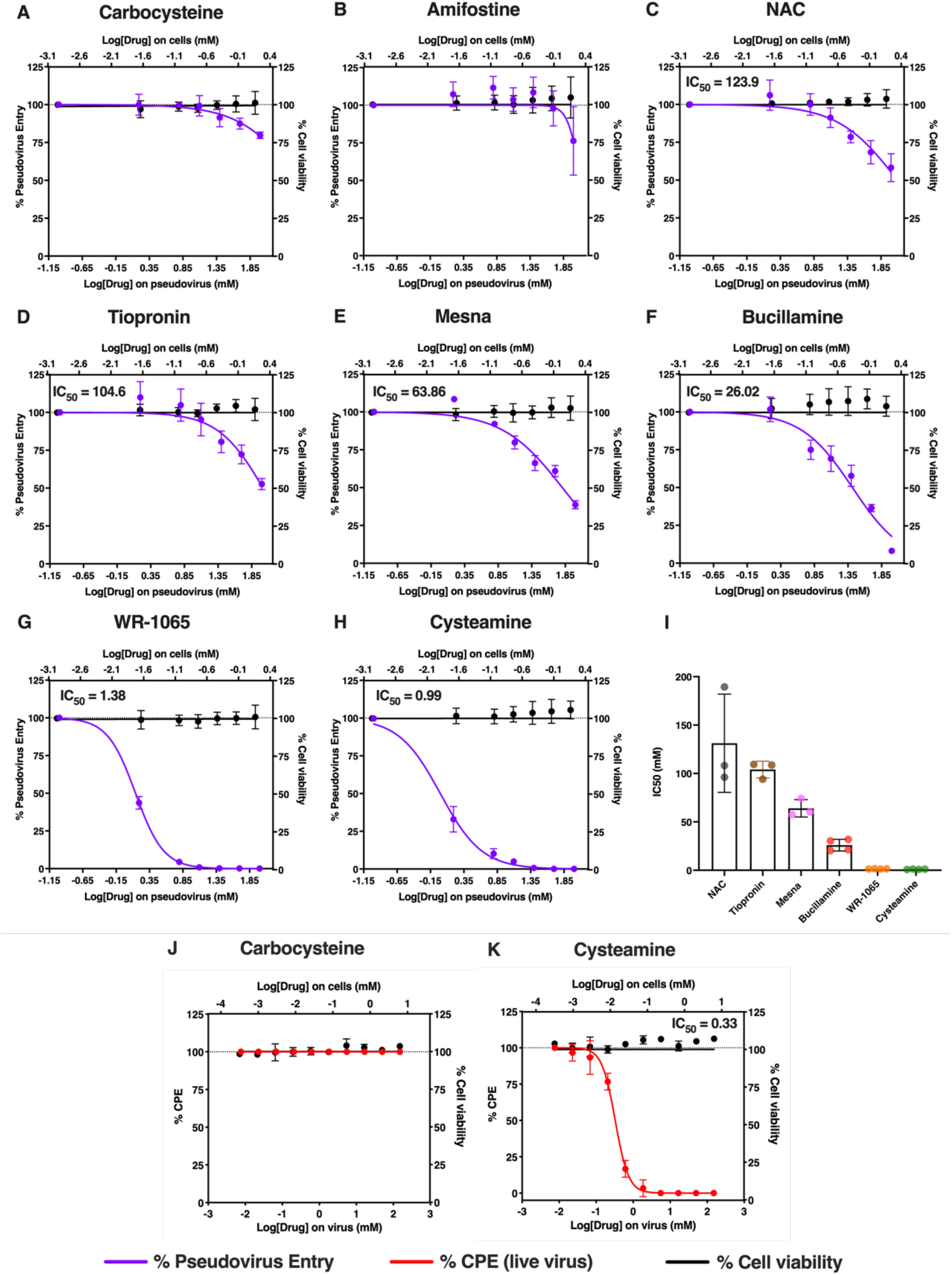
Thiol-based drugs inhibit SARS-CoV-2 in vitro. (**A to H**) Pseudovirus (PV) entry efficiency after pseudovirus exposure to (A) carbocysteine (sulfide drug, negative control), (B) amifostine (parent drug of WR-1065, negative control), thiol-based drugs (C) NAC, (D) Tiopronin, (E) Mesna, (F) Bucillamine, (G) WR-1065, and (H) Cysteamine prior to transduction into 293T-ACE2-TMPRSS2 cells (n = 3-4). The effects of drugs on cell viability were quantified using Cell Titer Glo 2.0 with lower drug dose exposures, reflecting 66-fold dilution of drugs when pseudovirus/drug mixture was incubated with cells (n=3). **(I)** IC50 values of the different thiol drugs in the PV assay. (**J-K**) Cytopathic effects (CPE) quantified by visual inspection when authentic SARS-CoV-2 is exposed to (J) carbocysteine (negative control) and (K) cysteamine (thiol drug) prior to infection in Vero E6-TMPRSS2 cells (n = 3). The effects of drugs on cell viability were quantified with exposure of cells to lower drug doses, reflecting the 24-fold dilution of drugs when virus/drug mixture was incubated with cells (n=3). The X-axes are scaled to log10 - the lower X-axis refers to the concentration of drugs on the PV/ live virus and the upper X-axis refers to concentration of drugs on the cells post dilution. Percentage changes are with respect to no drug control which is set as 100%. IC50 of the drugs was determined using the non-linear regression fitting with a variable slope. Data are mean ± SD.

### Thiol drugs inhibit entry mediated by SARS-2-S from multiple variants

Since the Delta variant (B.1.617.2) is now globally dominant, we tested the most potent thiol drugs in SARS-2-PV assays with spikes from B.1.617.2 (Delta variant) and the related B.1.617.1 (Kappa variant). In addition to Mesna, bucillamine, cysteamine and WR-1065, we included a novel thiol saccharide compound (methyl 6-**t**hio-6-**d**eoxy-*α*-D-**g**alactopyranoside [TDG]) previously proposed by us as a novel mucolytic because of its favorable properties as an inhaled drug (*45*). We found that all five drugs inhibited the entry of B.1.617.1 PV (Fig. 4A-F) and cysteamine and WR-1065 were the most potent (Fig. 4G). Pretreatment of cells followed by transduction with untreated PV had no significant effect (fig. S5A-F). In contrast to findings for B.1.617.1, the IC50 values for thiol drugs to inhibit B.1.617.2 variant entry were higher (Fig 4A-F). Whereas the IC50 value for bucillamine only increased by approximately two-fold, the IC50 values for TDG, Cysteamine and WR-1065 increased by approximately 3-15 fold and Mesna did not achieve an IC50 value at the doses tested (Fig. 4B-D). The IC50 values for Cysteamine and WR-1065 remained in the lower millimolar range (14mM and 6mM respectively) (Fig. 4E, F) and were the most potent of the thiol drugs tested (Fig 4G). Because cysteamine was consistently potent in inhibiting PV entry for Wuhan-1 and B.1.617 variants, we also tested its ability to inhibit entry of PVs for other variants (D614G, P.1., B 1.1.7, B.1.429 and B1.351). As shown in Fig. 4H-J, cysteamine was most potent against the Wuhan-1, Wuhan-1-D614G and B.1.617.1 variants but less potent against the other variants tested. The entry of the SARS-CoV-2 PV variants was not affected when the cells were pretreated with cysteamine prior to transduction with untreated PV (Fig S5 G, H).

**Fig. 4.**
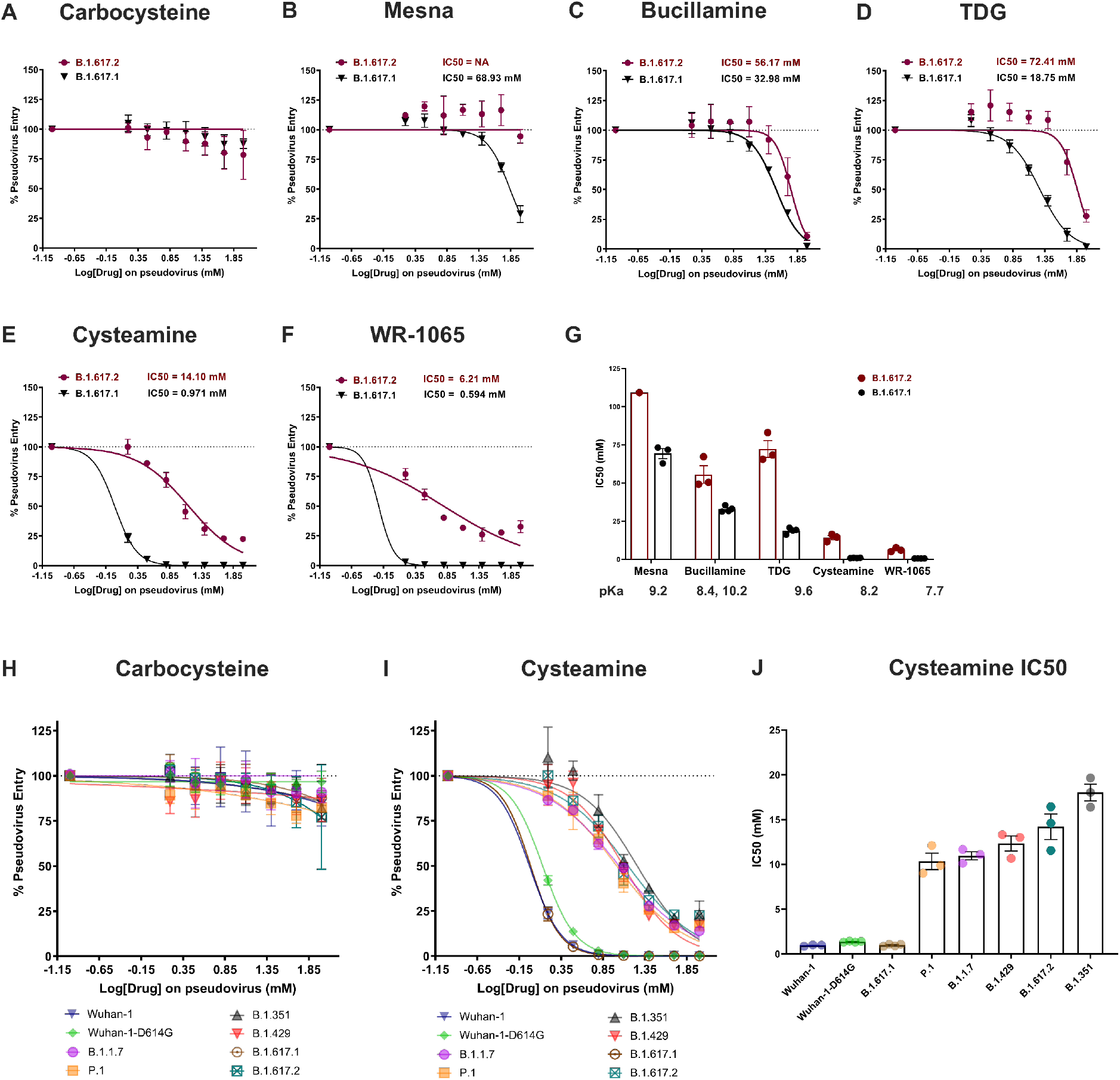
Thiol drugs reduce the infection of 293T-ACE2-TMPRSS2 cells by SARS-CoV-2 variant pseudoviruses. **(A to F)** Pseudovirus (PV) entry efficiency when the B.1.617.1 and B.1.617.1 pseudoviruses were exposed to (A) carbocysteine (sulfide drug, negative control), thiol drugs (B) Mesna, (C) Bucillamine, (D) TDG, (E) Cysteamine and (F) WR-1065 prior to transduction into 293T-ACE2-TMPRSS2 cells (n = 3-4). **(G)** IC50 values of the thiol drugs in the B.1.617.1 and B.1.617.2 PV transduction assay and their respective thiol pKa values. **(H, I)** Pseudovirus (PV) entry efficiency when the SARS-CoV-2 variant pseudoviruses were exposed to (H) carbocysteine (sulfide drug, negative control) and thiol drug (I) cysteamine prior to transduction into 293T-ACE2-TMPRSS2 cells (n = 3-4). **(J)** IC50 values of cysteamine in the SARS-CoV-2 variant PV transduction assays. NA = Not achieved.

### pKa is a determinant of thiol drug efficacy in SARS-2-PV assays

We noted that the potency of the thiol drugs is inversely related to their thiol pKa values (Fig. 4G) suggesting that a key physicochemical property of an optimal thiol drug to inhibit SARS-CoV-2 entry is a lower thiol pKa that is approaching the physiological pH. To further build support for this concept, we examined the efficacy of cysteine derivatives with a low pKa (e.g. cysteine methyl ester [CME], pKa = 7)(*46*) and a high pKa (e.g. N-acetyl cysteine, pKa = 9.5) in the SARS-2-PV assay using the B.1.617.1 variant. We found that the IC50 for CME was 25-fold lower than the IC50 for NAC and that L-cysteine (pKa = 8.4)(*46*) had intermediate potency (Fig. 5A-C, fig. S6A-C). These data indicate that the mechanism of action of thiol drugs as SARS-CoV-2 inhibitors is cleavage of disulfide bridges in SARS-2-S. To demonstrate that the cystine cleaving potency of these different thiol drugs in indeed related to their thiol pKa, we leveraged the BODIPY FL L-cystine reagent which fluoresces when thiol-specific exchange leads to mixed disulfide formation (Fig. 5D). We found that the cystine cleavage rates of thiol drugs in the BODIPY assay were inversely related to their pKa values (Fig. 5E, F) and mirrored the potency order seen in the SARS-2-PV assays. An outlier in this is TDG, a thiol-modified sugar with reported pKa of 9.6 but 5-fold higher potency than compounds with similarly high pKa values (NAC, Mesna). This finding indicates that pKa is not the only property that determines potency of a thiol drug; interactions between the thiol and RBD interface, hydrophilic or hydrophobic variations in each cystine microenvironment or steric factors may also affect potency.

**Fig. 5.**
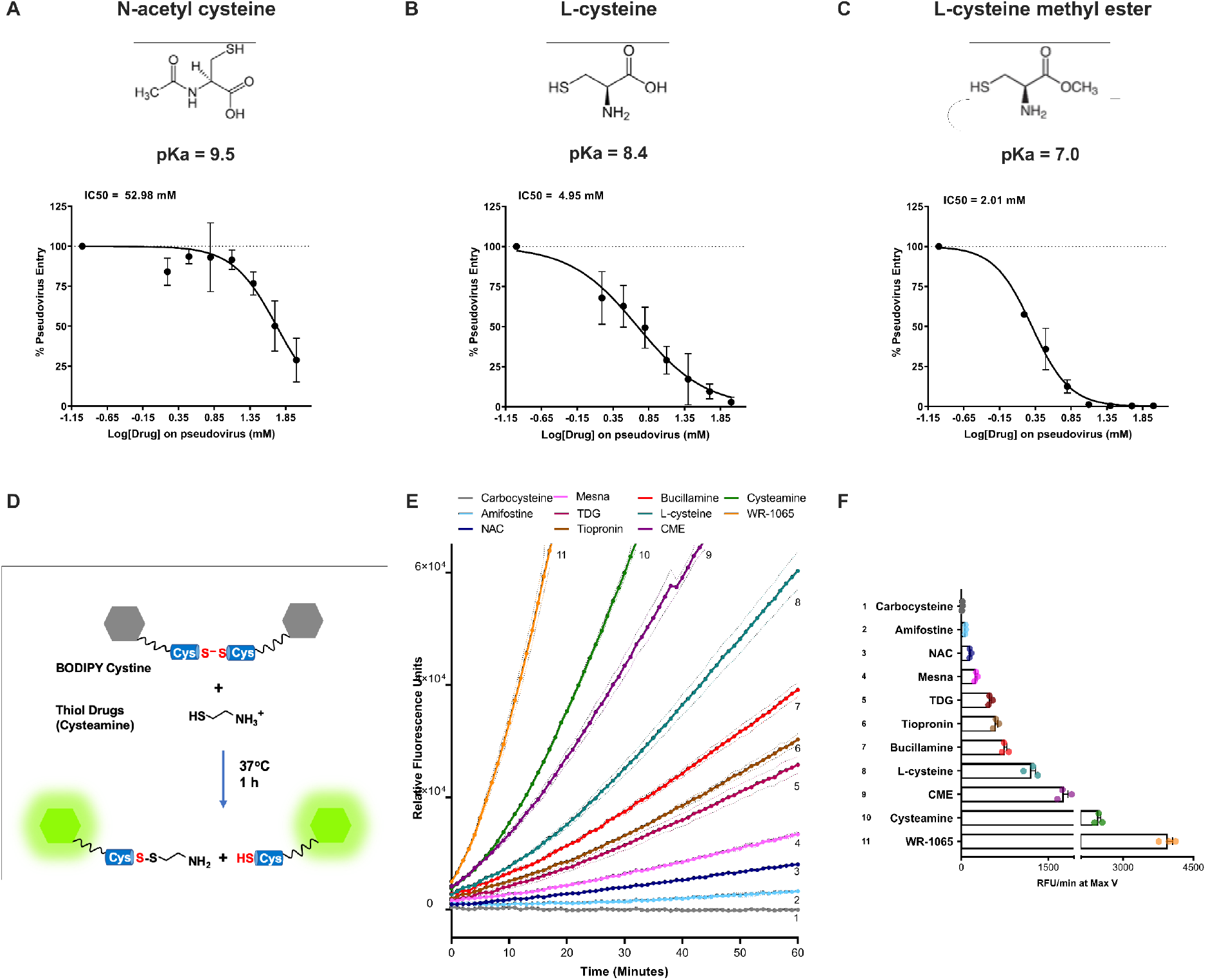
Thiol pKa influences the cystine cleaving rates of the thiol drugs and their efficacy in B.1.617.1 PV transduction assay. **(A to C)** PV entry efficiency when B.1.617.1 pseudoviruses were treated with three cysteine derivates (A) N-acetyl cysteine, (B) L-cysteine and (C) L-cysteine methyl ester (CME) having variable thiol pKa (n=3-6). Data are mean ± SD. (D) Schematic representation of the BODIPY cystine assay. (E) Rate of BODIPY FL-cystine cleavage when exposed to thiol drugs or controls (carbocysteine, amifostine) (n=3). Dotted lines indicate SEM. (F) Maximum slope (Max V) for the reactivity of thiol drugs with the BODIPY FL cystine reagent [based on data in (E)]. Numbers on graph (E) correspond with numbers designated to thiol compounds in (F). Data in (F) are mean ± SEM.

### Cysteamine reduces lung inflammation in a Syrian hamster model of COVID-19

We next tested if cysteamine reduces lung injury in the hamster model of SARS CoV-2 infection. We chose cysteamine as the thiol drug to test in these *in vivo* experiments for two reasons. First, cysteamine has a broad range of anti-oxidant anti-inflammatory activities (*11, 12, 19*), and it reduces aeroallergen-induced lung inflammation in mice (*20, 21*). Second, cysteamine was among the most potent thiol drugs in our *in vitro* antiviral assays described above. Cysteamine was administered intraperitoneally (IP) at a dose of 100 mg/kg to approximate the doses used clinically to treat cystinosis (*47, 48*)(Fig 6A). The first dose was given 2 hours prior to intranasal viral inoculation of SARS-CoV-2 (Wuhan-1, 2.94E+04 TCID_50_/animal) with subsequent doses given twice daily for 5 days (Fig. 6A). Outcomes of lung inflammation and lung injury in the hamsters were measured in bronchoalveolar lavage (BAL) fluid and in lung tissue. We found that multiple measures of lung inflammation were lower in cysteamine treated animals. Lung weights in cysteamine-treated hamsters were significantly lower than in the control group (Fig. 6C), and total protein in BAL were also lower (Fig. 6D). Total cell counts in BAL from cysteamine treated animals were also lower than in control treated animals with effects driven by decreases in neutrophils and lymphocytes (Fig. 6E-I). In addition, histopathology scores for mixed cell lung inflammation, composed primarily of macrophages and neutrophils, in lung tissue from cysteamine-treated animals were significantly lower than in the control group (Fig. 6J). Furthermore, alveolar hemorrhage scores from cysteamine-treated animals were significantly lower than in the control group (Fig. 6K, L). In contrast, viral N2 gene transcripts in lung tissue from cysteamine-treated animals were not significantly lower than in controls (Fig. 6B).

**Figure 6:**
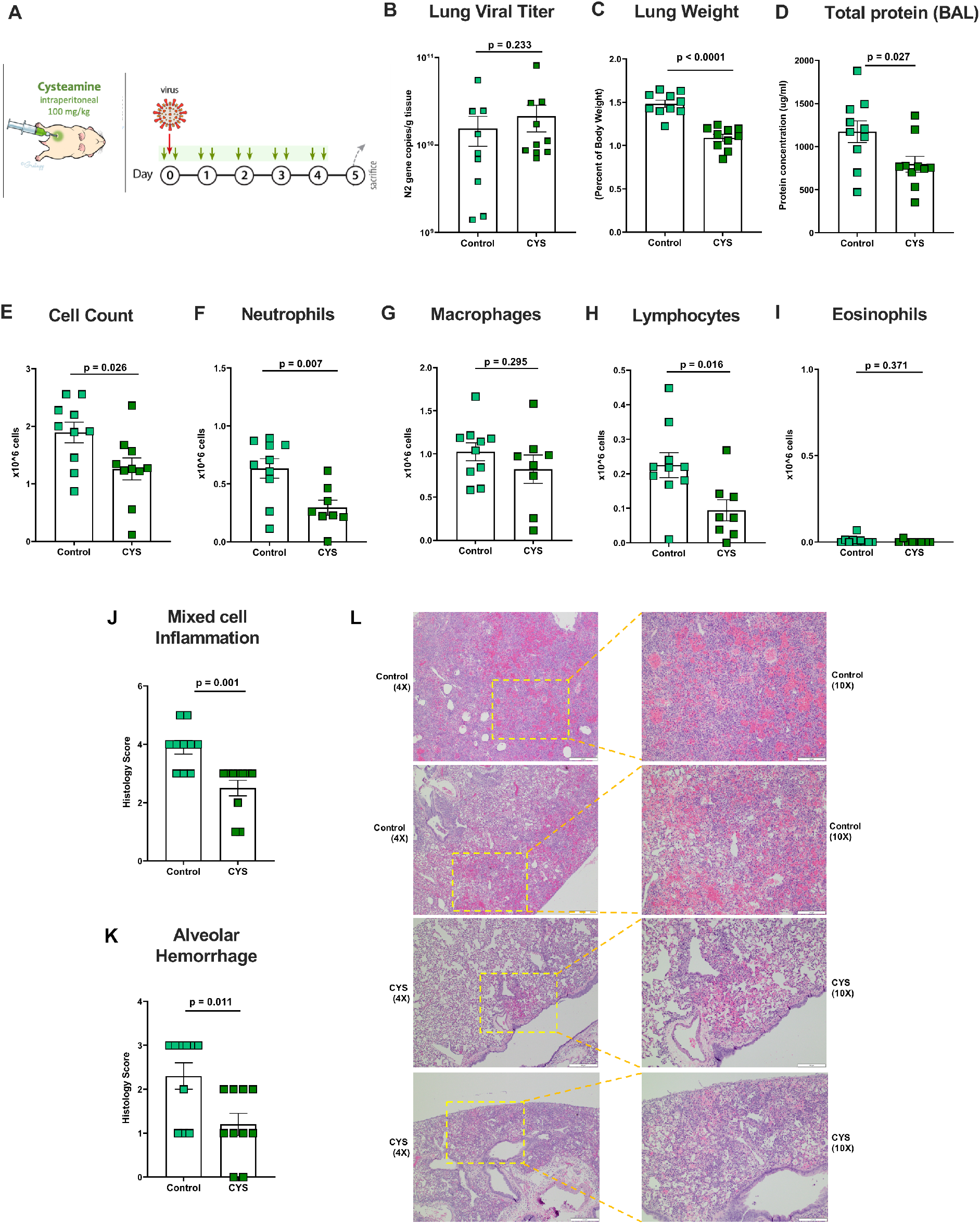
Effects of cysteamine on a Syrian hamster model of SARS-CoV-2 infection. (**A**) Study design for assessing the effect of cysteamine in Syrian hamster model of COVID-19. Cysteamine (100mg/kg) was administered twice daily via intraperitoneal injection for 5 days, with the first dose given 2 hours prior to the virus inoculation on Day 0. (**B**) Viral RNA levels in the lungs of animals treated with cysteamine relative to the vehicle control group. (**C**) Lung weights, normalized to the terminal body weights, of the animals. Total protein **(D)** and total cell counts **(E)** in the BAL fluid of hamsters treated with cysteamine with respect to the vehicle controls. (**F-I**) Differential leukocyte counts in the BAL fluid of animals, showing (F) neutrophil, (G) macrophage, (H) lymphocyte and (I) eosinophil counts in treated and vehicle control groups. (**J, K**) Histopathology scores for mixed cell (macrophages and neutrophils) inflammation in the peribronchovascular and the centriacinar regions of the lung (J) and alveolar hemorrhage (K) in the lungs of animals treated with cysteamine relative to vehicle control group. **(L)** Representative images of the lung sections of the animals highlighting the extent of alveolar hemorrhage in the animals from the two groups. Scale bar for 4X images equal 200μm; scale bar for 10X images equal 500μm. Control – intraperitoneal vehicle control group; CYS - cysteamine. Each group had N=10 animals. 2 BAL samples from the cysteamine group could not be analyzed due to a technical error. Data are mean ± SEM. Statistical significance was analyzed by two tailed, unpaired t-test between cysteamine treated (CYS) and respective control group (control).

## DISCUSSION

We explored if thiol drugs have anti-inflammatory and/or antiviral effects that could be beneficial in treating SARS-CoV-2 pneumonia. Our *in vivo* data in hamsters infected with SARS-CoV-2 shows that intra-peritoneal delivery of cysteamine significantly reduces lung inflammation and lung hemorrhage independent of any anti-viral effects. Our *in vitro* data in pseudovirus and live virus infection assays show that multiple thiol drugs cleave critical cystines in the spike protein of SARS-CoV-2 to prevent virus entry. The most potent thiol drugs have IC50 values in the low millimolar dose range, and these drugs concentrations are unlikely to be achieved in the airways by oral or systemic drug delivery. Thus, although thiol drugs have beneficial anti-inflammatory activity in SARS-CoV-2 pneumonia *in vivo* in hamster models, any antiviral activity *in vivo* in hamsters or in humans will require direct delivery to the airways to achieve needed drug concentrations in the lungs.

Cysteamine reduced SARS-CoV-2-related lung injury *in vivo* in hamsters without decreasing SARS-CoV-2 viral loads in the lung. Multiple outcomes of pneumonia improved in the cysteamine-treated animals, including decreases in lung neutrophils, total protein, and alveolar hemorrhage. Although we did not investigate the specific anti-inflammatory mechanisms for cysteamine efficacy in the SARS-CoV-2-infected hamsters, the positive therapeutic effects are plausibly explained by its anti-oxidant and anti-inflammatory properties. For example, cysteamine and other thiol drugs scavenge reactive oxygen species (ROS) and interrupt ROS-mediated inflammatory cascades, including pathways for NFkB activation and cytokine and chemokine production (*10–15*). Cysteamine can also replenish intracellular glutathione (*49–51*) and in its disulfide linked dimer form, it inhibits transglutaminase 2 (TG2) by thiol-disulfide interchange reactions (*18*). This is relevant because TG2 catalyzes transamidation reactions and crosslinks proteins to promote NFkB-dependent cytokine secretion (*20, 21, 52*). In addition, cysteamine inhibits somatostatin to limit substance P release and dampen neurogenic inflammation (*19*). The anti-inflammatory effects of cysteamine in the hamster COVID-19 model provides strong rationale to test it as an anti-inflammatory drug for COVID-19 in humans. Cysteamine is used most frequently as a drug to treat cystinosis, a lysosomal storage disease characterized by cystine accumulation, and it is approved in doses as high a 2g daily for life in these patients. Although it is not available as an intravenous formulation, it is available in tablet (including extended release) and eye drop formulations (*53, 54*).

The beneficial anti-inflammatory effects of cysteamine in COVID-19 may extend to the other 10 thiol drugs that are used as medicines (Table 1). All thiol drugs scavenge reactive oxygen species (ROS) and inhibit ROS-mediated inflammation in redox reactions that occur independently of their abilities to cleave cystine. Their anti-inflammatory actions are related to ROS scavenging, but a subset of thiol drugs have additional anti-oxidant or anti-inflammatory effects related to their ability to replenish glutathione directly (NAC, as a precursor) or indirectly (cysteamine, bucillamine, tiopronin) or ability to inhibit pro-inflammatory mediators (*49, 55–58*). For example, Mesna, tiopronin and D-penicillamine inhibit myeloperoxidase (*16, 17*) and, as noted above, cysteamine inhibits TG2 and somatostatin. Multiple thiol drugs including NAC, bucillamine, Mesna, penicillamine, tiopronin, erdosteine, and succimer are available in tablet form and can be taken orally (Table 1). Orally administered NAC is currently being tested in clinical trials in COVID-19 (NCT04419025, NCT04374461, NCT04703036) as is bucillamine, a thiol drug with two thiol warheads and efficacy in rheumatoid arthritis (*59, 60*)(NCT04504734). A feature of all thiol drugs is that they are generally well tolerated in high doses (up to multiple grams of drug per day) and a subset of thiol drugs can be administered intravenously to ensure high drug levels in blood. Intravenous delivery may therefore be a better clinical strategy for severe COVID-19 disease. NAC is used as a treatment for acetaminophen toxicity (*61*), where the dosing regimen involves intravenous administration of 20g of drug per day. Drugs other than NAC that are available as intravenous formulations include Mesna, amifostine and glutathione (Table 1). Amifostine is the only prodrug among these drugs - it is dephosphorylated to yield its active metabolite (WR-1065).

In addition to anti-oxidant and anti-inflammatory effects of thiol drugs, the ability of thiol drugs to cleave cystine confers these drugs with antiviral properties against SARS-CoV-2. We specifically hypothesized that thiol drugs cleave critical cystines in the RBD of SARS-2-S to disrupt binding to ACE2. In support of this hypothesis, we show in ACE2 binding assay, PV and live infection assays that multiple thiol drugs decrease binding of SARS-CoV-2 spike protein to its receptor to decrease virus entry into target cells. The potency of thiol drugs as viral entry inhibitors was highest for the Wuhan-1, Wuhan-1-D614G and B.1.617.1 (Kappa) variants where the most potent drugs - cysteamine and WR-1065 - had IC50 values in the 1 mM range. In contrast, the IC50 values for these drugs against the B.1.617.2 (Delta) variant were in the 5-15mM range. The Delta variant differs from the Kappa variant in having a lysine substitution (T478K) in its RBD (Fig 1) and the positive charge imparted by the amine groups in cysteamine and WR-1065 could mean that a charge-charge interaction impairs the access of these drugs to the RBD of the B.1.617.2 variant. While charge is a plausible explanation for differences in cysteamine potency among variants, it does not explain the higher IC50 values we found for thiol drug efficacy against variants with neutral amino acid substitutions (e.g. B.1.1.7 N501Y). In general, though, the conservation of cystines in the spike protein of SARS-CoV-2 VOIs and VOCs means that these drugs have potential utility as pan-variant therapeutics for COVID-19.

Low thiol pKa leading to high cystine cleaving activity was the key physicochemical property of the most potent thiol drugs. The ability of thiol drugs to cleave cystines is a function of the amount of drug in the deprotonated (thiolate) form. A pKa that approaches 7.4 results in higher fraction of active thiolate anion to act as a nucleophile in thiol-disulfide exchange reactions (*37, 62*). Cysteamine and WR-1065 have relatively low pKa values (8.2 and 7.7, respectively), and the relatively low pKa of cysteine methyl ester (pKa of 7.0) is the most likely explanation for its ability to inhibit the entry efficiency of SARS-CoV-2 more potently than N-acetyl cysteine (thiol pKa of 9.5). Together, these data reveal the importance of pKa as a key physicochemical property of thiol drugs for their efficacy as SARS-CoV-2 inhibitors based on cystine cleavage mechanism. Notably similar inverse dependence on the pKa related to thiolate concentration has been observed in reaction rates of thiol compounds with hydrogen peroxide and superoxide (*63*), suggesting that both anti-viral and anti-oxidant activity of thiols can be enhanced by selection of a thiol drug with low pKa.

The *in vitro* IC50 values for cysteamine is in the millimolar range, but systemic delivery of cysteamine is unlikely to deliver such millimolar drug concentration to the upper airways where SARS-CoV-2 first gains entry. For example, a 15 mg/kg dose of orally ingested cysteamine bitartrate resulted in peak plasma concentrations between 0.03 and 0.07 mM in children(*64*). Dose of 120 mg/kg dose cysteamine hydrochloride given IP to mice resulted in peak plasma concentration of 0.6 mM within 5 minutes after administration, followed by rapid decline (*65*). Thus, IP delivery of cysteamine is very unlikely to achieve millimolar concentrations in epithelial cells in the upper airways where the SARS-CoV-2 virus replicates and where viral titers are highest in early stages of infection (*66, 67*). For this reason, direct administration of cysteamine to the upper or lower airways via nasal spray or aerosol delivery is optimal to achieve the necessary millimolar drug concentrations in the airways and lungs. Other antiviral drugs, including remdesivir, are being formulated for inhaled route of administration in COVID19 (*67–69*), and a phase 2 clinical trial with inhaled nebulized interferon β-1a (IFNβ -1a) has shown promise in SARS-CoV-2 infection (*70*). Only a subset of currently approved thiol drugs (Mesna and NAC) are available as high dose liquid formulations that can be delivered by nebulizer (Table 1), and these drugs have high pKa values (> 9) and were among the least potent thiol drugs *in vitro* inhibition assays. The intermediate potency we found for methyl 6-**t**hio-6-**d**eoxy-a-D-**g**alactopyranoside [TDG] as a viral entry inhibitor for SARS-CoV-2 is noteworthy. Previously proposed as a novel mucolytic because of its favorable properties as an inhaled drug (*45*), it may be possible to design thiol-saccharide drugs with lower thiol pKa values than TDG. The design of such thiol-saccharide drugs can be informed by the chemical structures of cysteamine and WR-1065 which have positively charged amine groups close to their thiol warhead. Strategies such as introduction of electron-withdrawing groups to the saccharide scaffold to reduce thiol pKa can be used to synthesize compounds with potency similar or better than cysteamine and better suited to inhaled delivery.

In summary, thiol drugs reduce SARS-CoV-2-related lung inflammation and injury *in vivo*, and these data provide strong rationale for clinical trials of systemically delivered thiol drugs as treatments for SARS-CoV-2 pneumonia. Thiol drugs with lower thiol pKa values inhibit SARS-CoV-2 infection *in vitro* but in doses that will require drug delivery directly to the airways. Together our data reveal the potential for thiol drugs to act as anti-inflammatory and antiviral treatments for COVID-19, provide guidance for clinical trial designs of existing thiol drugs for treatment of COVID-19, and inform the design of and route of delivery for novel thiol drugs as antiviral treatments for COVID-19.

## MATERIALS AND METHODS

### Study Design

The purpose of this study was to investigate the antiviral and anti-inflammatory effects of thiol drugs in COVID-19. We used a plate-based colorimetric assay to test the effect of thiol drugs on binding of SARS-CoV-2 RBD to ACE2. Pseudovirus entry inhibition assays and authentic SARS-CoV-2 infection studies were performed to assess the antiviral effects of thiol drugs *in vitro*. All experiments were performed as at least three independent experiments with multiple replicates, as indicated in each section below. The *in vivo* efficacy of cysteamine was tested in a Syrian hamster model of SARS-CoV-2 infection. The study was conducted at the Lovelace Respiratory Research Institute (LBRI) using 20 hamsters (10 males, 10 females) with the sample size based on the published literature (*23, 24*) and the experience of the LBRI investigators in conducting drug efficacy studies for COVID19 in hamsters. Animals were placed in groups using Provantis with stratified (body weight) randomization so that animals with similar body weights were included in the study. Efficacy was determined by cell differentials from bronchoalveolar lavage (BAL) fluid, lung inflammation measured by lung weight gain and histopathology, and viral burden measured using RT-qPCR. Blinding to groups was not possible for this study. No data were excluded; 2 BAL samples from cysteamine treated group were not analyzed due to a technical error in collecting the BAL.

### Cells, plasmids and virus

HEK293T/clone17 (CRL-11268) were cultured in Dulbecco’s Modified Eaglés Medium (DMEM) supplemented with 10% fetal bovine serum and 1% penicillin/streptomycin (Thermo Fischer Scientific). The cells were obtained from ATCC and incubated at 37°C and 5% CO_2_. Vero cells stably over-expressing TMPRSS2 (Vero E6-TMPRSS2) (provided courtesy of Stefan Pölhmann (*27*)) were cultured in Dulbecco’s Modified Eaglés Medium (DMEM) supplemented with 10% fetal bovine serum and 1% penicillin/streptomycin (Thermo Fischer Scientific) and 5 μg/ml blastidicin (BSD; InvivoGen). MEXi 293E cells (IBA Lifesciences) were cultured in MEXi culture medium (IBA Lifesciences) at 37°C, 5% CO_2_ and 125 RPM as described by the manufacturer.

The codon-optimized SARS-CoV-2 spike gene was subcloned from pCG SARS-CoV-2 Spike (provided courtesy of Stefan Pölhmann (*27*)) into the EBNA-1 dependent expression vector pTT5 for high-level expression in MEXi 293E cells, while pCG SARS-CoV-2 B.1.617 (provided courtesy of Stefan Pölhmann (*71*)) was used to produce B.1.617 variant PVs. To boost cell surface expression of SARS-CoV-2 spike for efficient pseudotyping VSV, the C-terminal 21 amino acid containing the ER-retrieval signal (KxHxx) of spike was deleted. SARS-CoV-2, isolate USA-WA1/2020 (NR-52281) was obtained from BEI resources and passaged in Vero E6-TMPRSS2 cells (called as Wuhan-1 in text). Confluent Vero E6-TMPRSS2 cells grown in T175 flasks were infected with SARS-CoV-2 and the culture supernatant was collected when widespread cytopathic effect (CPE) was observed. After filtration through 0.45 μm filters, the virus containing culture supernatant was stored at −80°C in small aliquots.

### Thiol-based drugs and thiol content determination

N-acetylcysteine (NAC) and Mesna were used as the commercially available pharmaceutical formulations, with NAC manufactured by American Reagent INC at 200 mg/ml and Mesna by Baxter at 100 mg/ml USP. Cysteamine (MilliporeSigma), amifostine (MilliporeSigma), WR-1065 (MilliporeSigma) and penicillamine (MP Biomedicals) were provided as lyophilized powders that were solubilized as 500 mM concentrated stocks in water. Cysteamine and WR-2065 solutions were at pH 5. Amifostine solution was at pH 7 which was adjusted to pH 5 using 1M hydrochloric acid. To ensure that amifostine does not auto-dephosphorylate to WR-1065, the solution was made fresh before the experiment each time. Bucillamine (MilliporeSigma) and tiopronin (Spectrum Chemicals) were lyophilized powders that were solubilized as 500 mM concentrated stocks in equimolar NaOH to increase the solubility, and the pH was adjusted to pH 5. Carbocysteine (MilliporeSigma) and succimer (MilliporeSigma) were solubilized as 250 mM concentrated stocks in 500 mM NaOH to increase solubility with pH adjusted to pH 5. TDG (lyophilized powder synthesized according to (*45*)) was solubilized as 350mM stock in water. Free thiol content, and thus concentration of an active drug, was measured before every experiment using Ellman’s Reagent, 5,5’-dithio-*bis*-(2-nitrobenzoic acid) (DTNB) (Abcam), with the molar extinction coefficient of 14,150 M^-1^cm^-1^ at 412nm (*72*). Active drug concentration measured by DTNB was within 85 to 99% of nominal drug concentration. The stock solutions were stored at –20°C and discarded if the thiol content went below 85%. Drug concentrations reported in plate-binding and viral infection assays are based on active drug concentration in the stock solution.

### Structure Rendering and Analysis

Space filling images and receptor distance calculations were performed using indicated PDB entries with UCSF Chimera, developed by the Resource for Biocomputing, Visualization, and Informatics at the University of California, San Francisco, with support from NIH P41-GM103311 (*73*).

### RBD to ACE2 plate based binding assay

Wells of amine-reactive maleic anhydride-derivatized plates (Thermo Scientific) were coated overnight with recombinant SARS-CoV-2 Receptor binding domain (SARS-CoV-2 S protein RBD, His Tag (SPD-C52H3) containing amino acids Arg 319 - Lys 537 (Accession # QHD43416.1), ACRO Biosystems). The following day, the plates were washed and blocked with BSA for 1 hour at 37°C. Wells were then incubated for an hour at 37°C with drug solutions diluted in PBS at concentrations ranging from 0 to 20 mM. Negative controls included wells with no RBD or no ACE2. After washing, biotinylated soluble recombinant ACE2 (ACRO Biosystems) was added and incubated at 37°C for 60 minutes. After washing, streptavidin-HRP (ACRO Biosystems) was added to wells for an hour at 37°C. The plates were washed and incubated with TMB (Sera Care). The reaction was stopped and absorbance was read at 450 nm on a Biotek plate reader. Absorbance readings, after subtracting from negative control wells, were transformed to percent binding, with the wells containing no drug set as 100 percent binding. To measure the stability of binding of cysteamine, WR-1065, Mesna and bucillamine, wells were incubated with either of drugs at 5 mM for 1 hour, followed by three washes. ACE2 was then added to the wells either immediately, after 60 minutes, or after 120 minutes. Wells waiting for ACE2 were filled with dilution buffer. This was followed by the same steps to assess ACE2 binding as described above. For all binding assays, 4-6 independent experiments were carried out for all drugs.

### BODIPY FL L-cystine cleaving assay

BODIPY FL L-cystine (Thermo Fischer scientific) was reconstituted in methanol. In a black Maxi Sorp 96-well flat-bottomed plate (Nunc), 10 μM of BODIPY reagent prepared in PBS was mixed with 25 μM of the thiol-based drugs and the change in fluorescence was kinetically measured with excitation at 490nm and emission at 520nm, at 1minute intervals, for one hour at 37°C. Fluorescence reads, after subtracting the drug-free control reads, were plotted against time. The maximum slope (Max V) over a 10 minute interval for all the thiol-based drugs from this plot was calculated and represented as relative fluorescence units/min (RFU/min) to assess the cystine cleaving ability of the drugs. The experiment was repeated three times.

### Production of pseudoviruses

Pseudoviruses bearing SARS-2-S were generated as previously described using recombinant VSVΔG-luciferase-based viruses, which lack glycoprotein (G) gene and instead code for reporter gene firefly luciferase (*44*). Briefly, either MEXi (IBA LifeSciences) or BHK-21/WI-2 (Kerafast) cells were transfected with SARS-CoV-2 Spike expression plasmid (pTT5 SARS-CoV-2 SΔ21), using polyethylenimine (PEI) or TransIT-2020 (Mirus Bio) as described by the manufacturer. Mock transfection served as the ‘no glycoprotein’ control. At 24 hours post-transfection, the cells were inoculated with VSVΔG-luc(VSV-G) at a multiplicity of infection (MOI) of 0.3-3. After 6 hours of incubation, the cells were washed three times with PBS and resuspended in culture medium containing 1% I1 anti-VSV-G hybridoma supernatant (ATCC CRL-2700). At 24 hours post-infection, the culture supernatant was collected by centrifugation and filtered through a 0.45 μm syringe filter to clear off cellular debris. The supernatant containing viral particles was aliquoted and stored at −80 °C until further use.

### Establishment of HEK293T cells stably expressing ACE2 and TMPRSS2 (293T-ACE2-TMPRSS2)

ACE2 was cloned into pLKO5d.SFFV.dCas9-KRAB.P2A.BSD (a gift from Dirk Heckl, Addgene plasmid) and TMPRSS2 was cloned into pDUAL CLDN (GFP) (a gift from Joe Grove, Addgene plasmid). All cloning steps were confirmed by Sanger sequencing. Lentiviral particles for delivery of lentiviral ACE2 and TMPRSS2 vectors were produced using transfection with PEI. Briefly, HEK293T cells were transfected with three plasmids: lentiviral ACE2 or TMPRSS2 constructs, psPAX2, and pVSV-G. 48 h post transfection, cell supernatants containing the newly produced viral particles were centrifuged and subsequently filtered using 0.22 µm vacuum filter units (MilliporeSigma). The supernatants were then aliquoted and stored at −80°C. To establish 293T-ACE2-TMPRSS2 cells, HEK293T cells were transduced with lentiviral particles containing the ACE2 vector. 48h post transduction, medium was replaced with blastidicin (BSD; InvivoGen) selection medium. After 5 days of selection, further expansion of cells stably expressing ACE2 was carried out. The process was then repeated to further transduce cells with TMPRSS2 lentiviral particles and cells were cultured in antibiotic selection medium containing BSD and Puromycin 48h post transduction. The expression of ACE2 and TMPRSS2 was confirmed by Western Blot and compared to non-transduced cells.

### Pseudovirus transduction experiments

293T-ACE2-TMPRSS2 cells were plated in black 96-well tissue culture treated plates (Greiner Bio-one) 18 hours before the experiment. Two experimental strategies of pseudovirus pre-treatment and cell pre-treatment were followed (fig. S1). For pseudovirus pre-treatment, the pseudoviruses were pre-incubated with different concentrations (1.56-100mM) of the thiol-based drugs for 2 hours at 37°C, followed by 66-fold dilution with standard culture media. The cells were then transduced with these pre-treated virions for 2 hours at 37°C. After the incubation, the virions were removed and cells were cultured in standard culture medium. For cell re-treatment, the 293T-ACE2-TMPRSS2 cells were incubated with the different drug concentration (0.02 -1.5mM) for 2 hours at 37°C, 5%CO_2_. These concentrations reflect the 66-fold dilution of drugs when virus/drug mix was incubated with the cells in the pseudovirus pre-treatment experiment. After incubation, the media was removed and the cells were transduced with untreated pseudoviruses for 2 hours at 37°C. After the incubation, the virions were removed and the cells were cultured in standard culture medium. For both experimental conditions, at 18 hours post-transduction, the cells were lysed and luciferase activity was measured using Promega luciferase assay system and Biotek Synergy H1 plate reader. Data was normalized to the viral particles without any viral envelope protein. For each experiment, luciferase reads of drug free group was set as 100% and the relative transduction efficiencies in the presence of thiol-based drugs were calculated. Three-four independent experiments were carried out for each PV pretreatment and cell pretreatment strategies, with 6-12 replicates in each for all the drug doses.

### Inhibition of authentic SARS-CoV-2 (Wuhan-1) infection

SARS-CoV-2 of 1.5 x 10^3^ TCID_50_/ml was incubated with 3-fold serially diluted carbocysteine and cysteamine (0.007-150 mM) at 37°C for 2 hrs. Virus-drug mixtures were diluted 24-fold before addition to Vero E6-TMPRSS2 cell monolayer in 96-well-plate. For each drug concentration, virus-drug mixtures were added to 10 replicate wells at 100 μl per well. The final titer of virus added to cells was 63.2 TCID_50_/ml. After two hours of infection, virus-drug inoculum was replaced with fresh DMEM medium containing 2% FBS. Clear CPE developed after 7 days of incubation at 37°C with 5% CO_2_. The experiment was repeated thrice. Wells with clear CPE were counted positive and percentage of positive wells for each concentration of tested drugs were plotted. The effect of thiol-based drugs on Vero E6-TMPRSS2 cells during the two hours of SARS-CoV-2 infection was evaluated by addition of 6.25 mM of each drug and 63.2 TCID_50_/ml SARS-CoV-2 simultaneously to Vero E6-TMPRSS2 cell monolayer in 96-well-plate. This concentration reflects the highest concentration, post 24-fold dilution of drugs when virus/drug mix was incubated with the cells in the virus pre-treatment experiment. After two hours of infection, cells were washed and then cultured with fresh DMEM medium containing 2% FBS at 37°C with 5% CO_2_. Clear CPE developed 7 days post infection.

### Quantification of cell viability

The cell viability was quantified using CellTiter-Glo2.0 assay (Promega) which measures cellular ATP content, indicating the metabolically active cells. For all cell viability experiments, the experimental protocol was the same as the main experiment except for the step of pseudovirus/live virus infection. For cell viability measurement corresponding to pseudovirus experiment, 293T-ACE-TMPRSS2 cells were seeded in 96 well black plates 18 hours prior to the experiment. The cells were then incubated with different concentrations (0.02 -1.5mM) of the thiol-based drugs for 2 hours at 37°C, followed by removal of the drugs and incubation of cells with standard culture medium for 18 hours. The experiment was carried out thrice with 5-6 replicates for each drug. These concentrations reflect the 66-fold dilution of drugs when pseudovirus/drug mix was incubated with the cells in the pseudovirus pretreatment setting. For the cell viability measurement corresponding to the live virus experiment, Vero E6-TMPRSS2 cells were incubated with different concentrations (0.0003-6.25mM) of the drugs in 1% FBS for 7 days. These concentrations reflect the 24-fold dilution of drugs when virus/drug mix was incubated with the cells in the live virus infection setting. The cell viability experiment on Vero E6-TMPRSS2 cells was carried out thrice with 6 replicates for each drug. For both cell viability experiments, post the respective incubations, the plates and their contents were equilibrated at room temperature for 30 minutes before addition of equal volumes of CellTiter Glo2.0 reagent. Afterwards, the contents were mixed on a plate shaker to induce cell lysis. The plates were then incubated at room temperature for 10 minutes followed by measurement of luminescence using Biotek plate reader. Luciferase reads of control-treated cells was set as 100% and the relative viability of cells incubated in the presence of thiol-based drugs was calculated.

### Syrian Hamster Model of COVID-19

The efficacy of cysteamine was tested in a Syrian hamster model of SARS-CoV-2 infection. All the antiviral studies were performed in an animal biosafety level 3 (ABSL3) facility at the Lovelace Respiratory Research Institute, Albuquerque, New Mexico. All work was conducted under protocols approved by the Institutional Animal Care and Use Committee (IACUC). A total of 20 Syrian hamsters (*Mesocricetus auratus*), with a target age of 6-10 weeks old and a body weight of 115-136 g (mean± SD = 126.1 g ± 6 g), were on the study. The animals were divided into 2 groups – animals receiving intraperitoneal dosing of the vehicle (water), and animals receiving cysteamine hydrochloride (147 mg/kg; equal to 100mg/kg cysteamine free base; MilliporeSigma). Cysteamine is used in high doses clinically (2 g/day). A dose of 100 mg/kg was administered to hamsters, based on conversion of human daily dose (2 g/day) to animal dose based on body surface area using FDA guidance(*74*). The animals were dosed twice daily for 5 days, starting on Day 0, as shown in Fig. 6A. The first dose was administered 2 hours prior to the viral inoculation. All animals were inoculated intranasally with SARS-CoV-2 (isolate USA-WA1/2020 or Wuhan-1) at 2.94×10^4^ TCID_50_/animal. Efficacy of the drug was determined by measuring viral load by RT-qPCR and lung inflammation, as measured by lung weight gain, total and differential cell counts in the bronchoalveolar lavage fluid and histopathological analysis of the lung sections.

#### Bronchoalveolar Lavage (BAL) collection and processing

BAL was performed on the right lung lobe. The left lobe was clamped off and the right lung lobes were lavaged with sterile saline. Half of the BAL collected was UV irradiated for sterilization out of the ABSL3 and used for differential analysis. This aliquot was centrifuged at 1000 g, 2-8°C, ≥10 minutes. The cell pellet was resuspended in the appropriate amount of resuspension buffer, and red blood cell lysis buffer was used on samples as necessary. The total cell count was counted using a Nexcelom automated cell counter. A total of 50,000 cells per slide were used to prepare microscope slides by cytocentrifugation. The cells on slides were fixed and stained using Modified Wright’s or Wright-Giemsa Stain. Differential counts on at least 200 nucleated cells per slide were conducted using morphological criteria to classify cells into neutrophils, macrophages, lymphocytes and eosinophils.

#### Histopathology

At scheduled necropsy, animals were euthanized by intraperitoneal injection with an overdose of a barbiturate-based sedative. The right lobes of the lungs were used for BAL collection and processed to measure viral loads. The left lobes of the lungs were collected, examined, and weighed, and representative samples were preserved for histopathology. Lung lobes were instilled via major airway(s) with 10% neutral buffered formalin (NBF) to approximate physiologic full lung volume at 25cm hydrostatic pressure; the major airway(s) used for instillation were closed, and the lobes were immersed in NBF for fixation. Tissues were trimmed and processed routinely, paraffin embedded, sectioned at 4 μm and stained with hematoxylin and eosin for microscopic examination. Histopathologic examination was conducted on the left lung lobes with findings for a given tissue graded subjectively and semi-quantitatively by a single board-certified veterinary pathologist on a scale of 1-5 (1 = Minimal, 2 = Mild, 3 = Moderate, 4 = Marked, 5 = Severe). The Provantis™ v10.2.3.1 (Instem LSS Ltd., Staffordshire, England) computer software/database was used for necropsy and histopathology data acquisition, reporting and analysis.

#### Viral titers using RT-qPCR

Lung samples were homogenized in Trizol using a TissueLyser and centrifuged at 4000 x g for 5 minutes. From the supernatants, RNA was isolated using the Direct Zol-96 RNA Kit (Zymo Research), according to the manufacturer’s instructions.

SARS-CoV-2 viral RNA was quantified by a qPCR assay targeting the SARS-CoV-2 nucleocapsid phosphoprotein gene (N gene). Genome copies per g equivalents were calculated from a standard curve generated from RNA standards of known copy concentration. All samples were run in triplicate. The SARS-CoV-2 N gene primers and probe sequences are as follows:

SARS-CoV-2 Forward: 5’ TTACAAACATTGGCCGCAAA 3’

SARS-CoV-2 Reverse: 5’ GCGCGACATTCCGAAGAA 3’

SARS-CoV-2 Probe: 6FAM-ACAATTTGCCCCCAGCGCTTCAG-BHQ-1

Amplification and detection was performed using a suitable real-time thermal cycler under the following cycling conditions: 50 °C for 5 minutes, 95 °C for 20 seconds and 40 cycles of 95 °C for 3 seconds, and 60 °C for 30 seconds.

### Statistical analysis

Tests, number of animals (n), and statistical comparison groups are indicated in the Figure legends. For analyzing the statistical significance of difference in loss of binding for each drug area under the curve (AUC) was plotted and ordinary one -way ANOVA followed by Dunnett’s post hoc analysis was performed. Data are presented as mean ± SEM [** p ≤ 0.01, **** p ≤ 0.0001]. IC50 of the drugs in pseudovirus transduction and authentic SARS-CoV-2 infection experiments was determined using the non-linear regression fitting with a variable slope. Data for pseudovirus entry inhibition and authentic SARS-CoV-2 infection experiments are plotted as mean ± SD. For the *in vivo* data, statistical significance was analyzed by two tailed, unpaired t-test between cysteamine treated and the control groups. All statistical analyses were performed using GraphPad Prism software (version 9.2).

## Supplementary Materials

### Supplementary methods

#### Supplementary figures

Fig. S1. Effect of penicillamine and succimer on binding of SARS-CoV-2 RBD to ACE2.

Fig. S2. Schematic illustration of the strategies employed to assess thiol-based drugs as pseudovirus entry inhibitors.

Fig. S3. Effect of pretreatment of 293T-ACE2-TMPRSS2 cells with thiol drugs on entry of pseudoviruses (ancestral strain/ Wuhan-1) in the cells.

Fig. S4. Effect of thiol drugs on authentic SARS-CoV-2 infection (Wuhan-1).

Fig. S5. Effect of pretreatment of 293T-ACE2-TMPRSS2 cells with thiol drugs on entry of variant pseudoviruses.

Fig. S6. Effect of pretreatment of 293T-ACE2-TMPRSS2 cells with cysteine derivatives on B.1.617.1 pseudovirus entry.

## Acknowledgments

The authors would like to thank the technical staff at Lovelace Respiratory Research Institute for their work infecting hamsters with SARS-CoV-2 in the ABSL3 facility. The authors also thank Chris Gralapp for drawing Figures 2A and 6A.

The following reagent was deposited by the Centers for Disease Control and Prevention and obtained through BEI Resources, NIAID, NIH: SARS-Related Coronavirus 2, Isolate USA-WA1/2020,NR-52281.

## Author contributions

K.K.: Conceptualization; methodology; visualization; writing - original draft; writing - reviewing and editing.

W.R.: Conceptualization; methodology; visualization; writing - reviewing and editing.

A.R.C.: methodology; visualization; writing - reviewing and editing.

J.J.: methodology; writing - reviewing and editing.

I.G.: Conceptualization; writing - reviewing and editing.

M.T.: Methodology.

A.W.: Methodology.

T.B.: Methodology.

J.C.: Methodology.

S.B.: Methodology.

H.S.S.: Methodology.

S.F.: Methodology.

R.M.: Methodology.

S.Pillai: Writing - reviewing and editing.

A.M.H.: Methodology.

S.O.: Methodology.

M.H.: Methodology.

S.P.: Methodology, Writing - reviewing and editing.

G.S.: Supervision; methodology; writing - reviewing and editing.

J.V.F.: Conceptualization; supervision; methodology; visualization; writing - original draft; writing - reviewing and editing.

## Data and materials availability

All data are available in the main text or the supplementary materials.

