## Supplementary File for "Thiol drugs decrease SARS-CoV-2 lung injury *in vivo* and disrupt SARS-CoV-2 spike complex binding to ACE2 *in vitro*"

**This PDF file includes:**

Materials and Methods
Fig. S1 to S6

**Materials and Methods**

**Inhibition of authentic SARS-CoV-2 infection**

For the live virus infection data in the supplement section, SARS-CoV-2 of 1.2 x $10^4$ $TCID_{50}$/ml was incubated with 2-fold serially diluted thiol-based drugs (0.4-6.25 mM for cysteamine and WR-1065, 6.25-100mM for Mesna and bucillamine) at 37°C for 2 hrs. Virus-drug mixtures were diluted 12-fold before addition to Vero E6 cell monolayer in 96-well-plate. For each drug concentration, virus-drug mixtures were added to 10 replicate wells at 100 µl per well. The final titer of virus added to cells was 1 x $10^3$ $TCID_{50}$/ml (100 $TCID_{50}$ per 100 µl per well in 96-well-plate). After two hours of infection, virus-drug inoculum was replaced with fresh DMEM medium containing 1% FBS. Clear CPE developed after two days of incubation at 37˚C with 5% $CO_2$. The experiment was repeated thrice. Wells with clear CPE were counted positive and percentage of positive wells for each concentration of tested drugs were plotted. The effect of thiol-based drugs on Vero E6 cells during the two hours of SARS-CoV-2 infection was evaluated by addition of 8.33 mM or 0.52 mM of each drug and 100 $TCID_{50}$ SARS-CoV-2 simultaneously to Vero E6 cell monolayer in 96-well-plate. These concentrations reflect highest concentration, post 12-fold dilution of drugs when virus/drug mix was incubated with the cells in the virus pre-treatment experiment. After two hours of infection, cells were washed and then cultured with fresh DMEM medium containing 1% FBS at 37˚C with 5% $CO_2$. Clear CPE developed two days post infection.

**Quantification of cell viability**

The cell viability was quantified using CellTiter-Glo2.0 assay (Promega) which measures cellular ATP content, indicating the metabolically active cells. For the cell viability measurement corresponding to the live virus experiment, Vero E6 cells were incubated with different concentrations (0.03-8.3mM) of the drugs in 1% FBS for 7 days. These concentrations reflect the 12-fold dilution of drugs when virus/drug mix was incubated with the cells in the live virus infection setting. The cell viability experiment on Vero E6 cells was carried out thrice with 6 replicates for each drug. After the incubations, the plates and their contents were equilibrated at room temperature for 30 minutes before addition of equal volumes of CellTiter Glo2.0 reagent. Afterwards, the contents were mixed on a plate shaker to induce cell lysis. The plates were then incubated at room temperature for 10 minutes followed by measurement of luminescence using Biotek plate reader. Luciferase reads of control-treated cells was set as 100% and the relative viability of cells incubated in the presence of thiol-based drugs was calculated.

**Figure S1.**

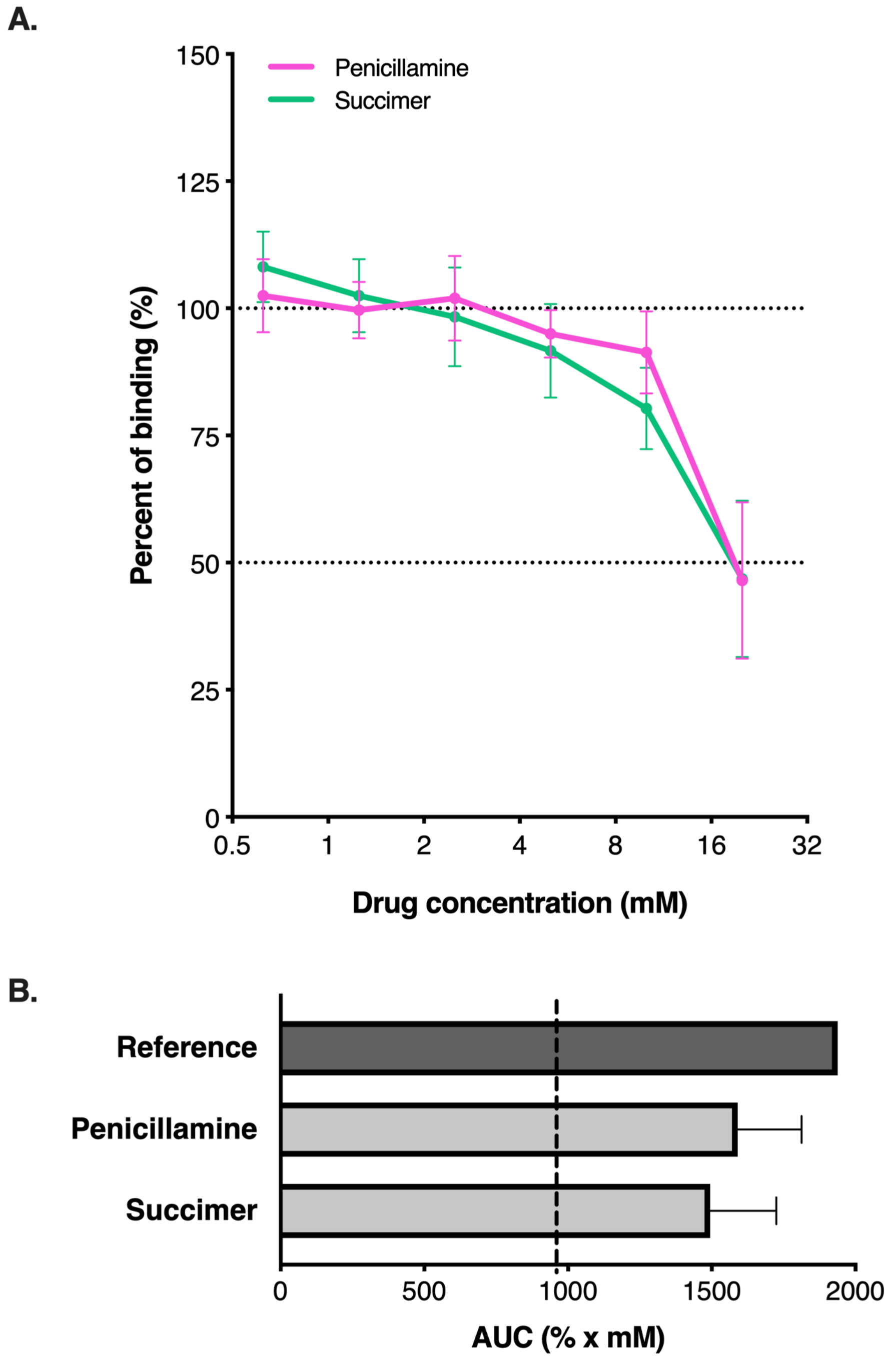


**Figure S1. Effect of penicillamine and succimer on binding of SARS-CoV-2-RBD to ACE2.**

**(A)** Percent of binding in the presence of the penicillamine and succimer (n = 4 - 6). Without drug treatment, the binding was 100%, whereas treatment with the thiol-based drugs showed a decrease in the binding % relative to no drug control. The X axis is scaled to log2. **(B)** Area under the curve (AUC) analysis for effects of the thiol-based drugs on RBD to ACE2 binding. Reference AUC was calculated from RBD to ACE2 binding with no drug control; dashed line represents 50% of reference AUC. Data are mean ± SEM. Statistical significance was analyzed by one-way ANOVA followed by Dunnett's post-hoc analysis. Significance indicates differences from reference AUC.

**Figure S2.**

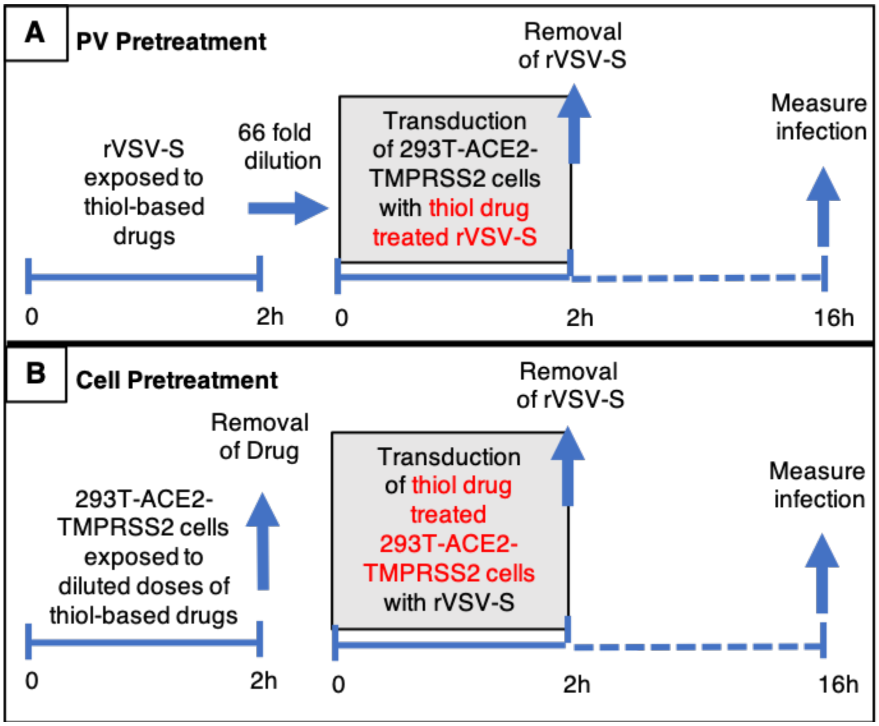


**Figure S2. Schematic illustration of the strategies employed to assess thiol drugs as pseudovirus entry inhibitors. (A)** Pseudovirus (PV) pretreatment: The rVSV-S was pre-incubated with thiol-based drugs prior to transduction of the 293T-ACE2-TMPRSS2 cells. **(B)** Cell pretreatment: The 293T-ACE2-TMPRSS2 cells were exposed to thiol-based drugs before the cells were transduced with rVSV-S.

**Figure S3.**

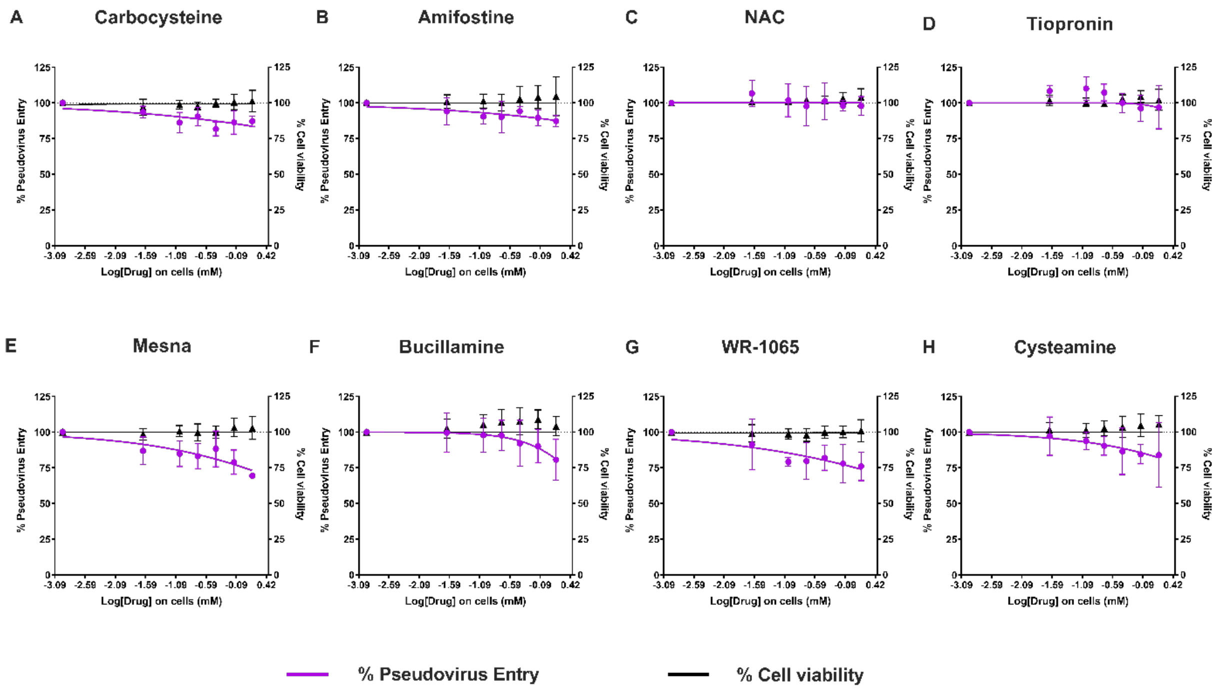


**Figure S3. Effect of pretreatment of 293T-ACE2-TMPRSS2 cells with thiol drugs on the entry of pseudoviruses (ancestral strain Wuhan-1) in the cells.** Pseudovirus entry efficiency, quantified by luciferase activity, when the cells were exposed to drugs prior to transduction with untreated virus (as illustrated in Figure S2B), (n = 3). The effects of drugs on viability of 293T-ACE2-TMPRSS2 cells was quantified using Cell Titer Glo 2.0 (n=3). The drug doses reflect the 66-fold dilution of drugs when pseudovirus/drug mixture was incubated with cells in the pseudovirus pretreatment strategy. The X-axis are scaled to log10. Percentage changes are with respect to no drug control which is set as 100%. Data are mean ± SD.

**Figure S4**

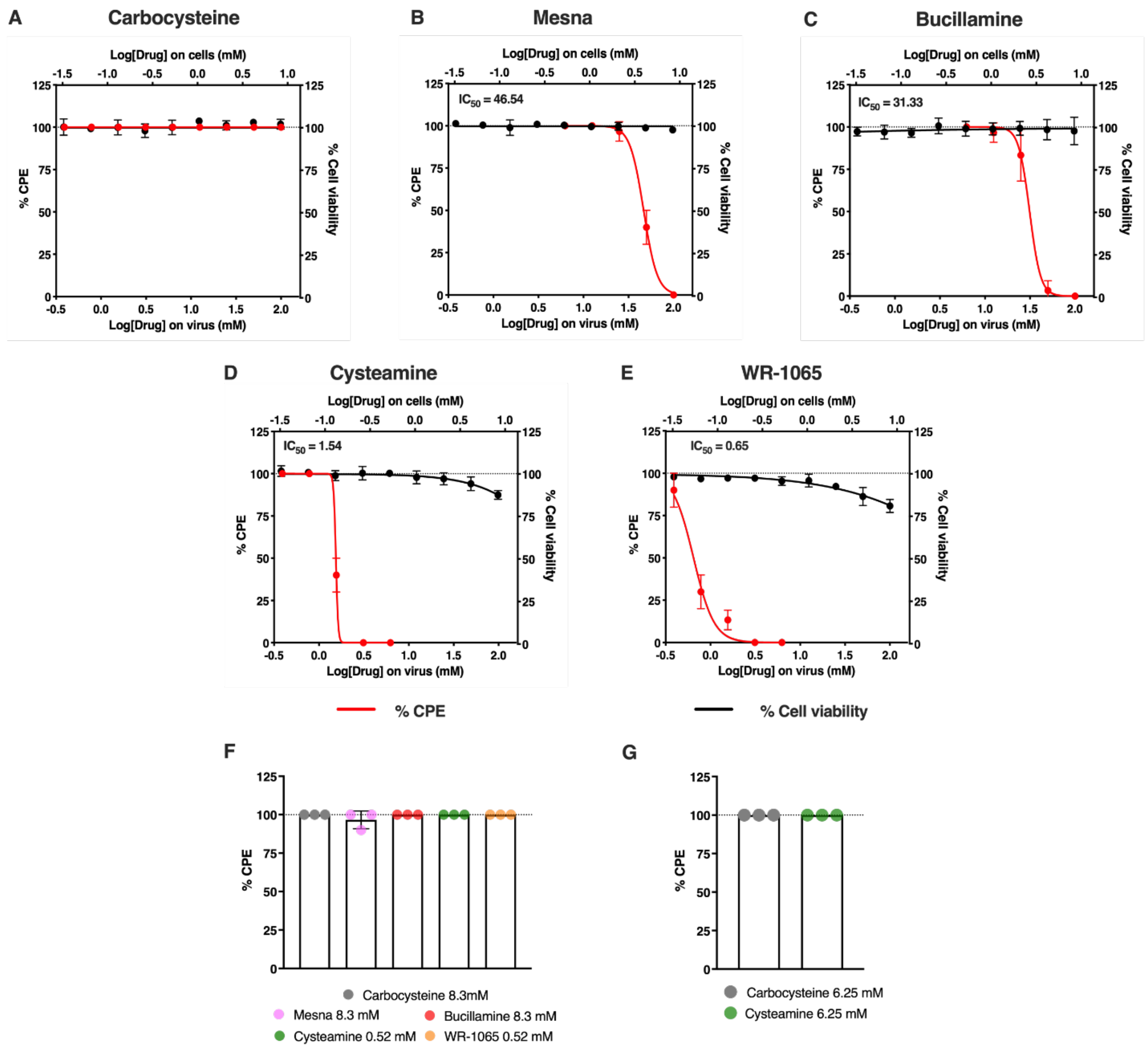


**Figure S4. Effect of thiol drugs on authentic SARS-CoV-2 infection (Wuhan-1). (A** to **E)** Cytopathic effects (CPE) quantified by visual inspection when virus is exposed to (A) carbocysteine (negative control) and thiol drugs (B) Mesna, (C) Bucillamine, (D) Cysteamine, (E) WR-1065 prior to infection in Vero E6 cells (n = 3). **(F)** The effects of drugs on Vero E6 cell viability were quantified with exposure of cells to lower drug doses, reflecting the 12-fold dilution of drugs when virus/drug mixture was incubated with cells (n=3). **(G)** The effects of drugs on

Vero-TMPRSS2 cell viability were quantified with exposure of cells to lower drug doses, reflecting the 24-fold dilution of drugs when virus/drug mixture was incubated with cells (corresponding to Fig. 3J,K) (n=3). The X-axes are scaled to log10 - the lower X-axis refers to the concentration of drugs on the live virus and the upper X-axis refers to equivalent concentration of drugs on the cells. Percentage changes are with respect to no drug control which is set as 100%. IC50 of the drugs was determined using the non-linear regression fitting with a variable slope. Data are mean ± SD.

**Figure S5.**

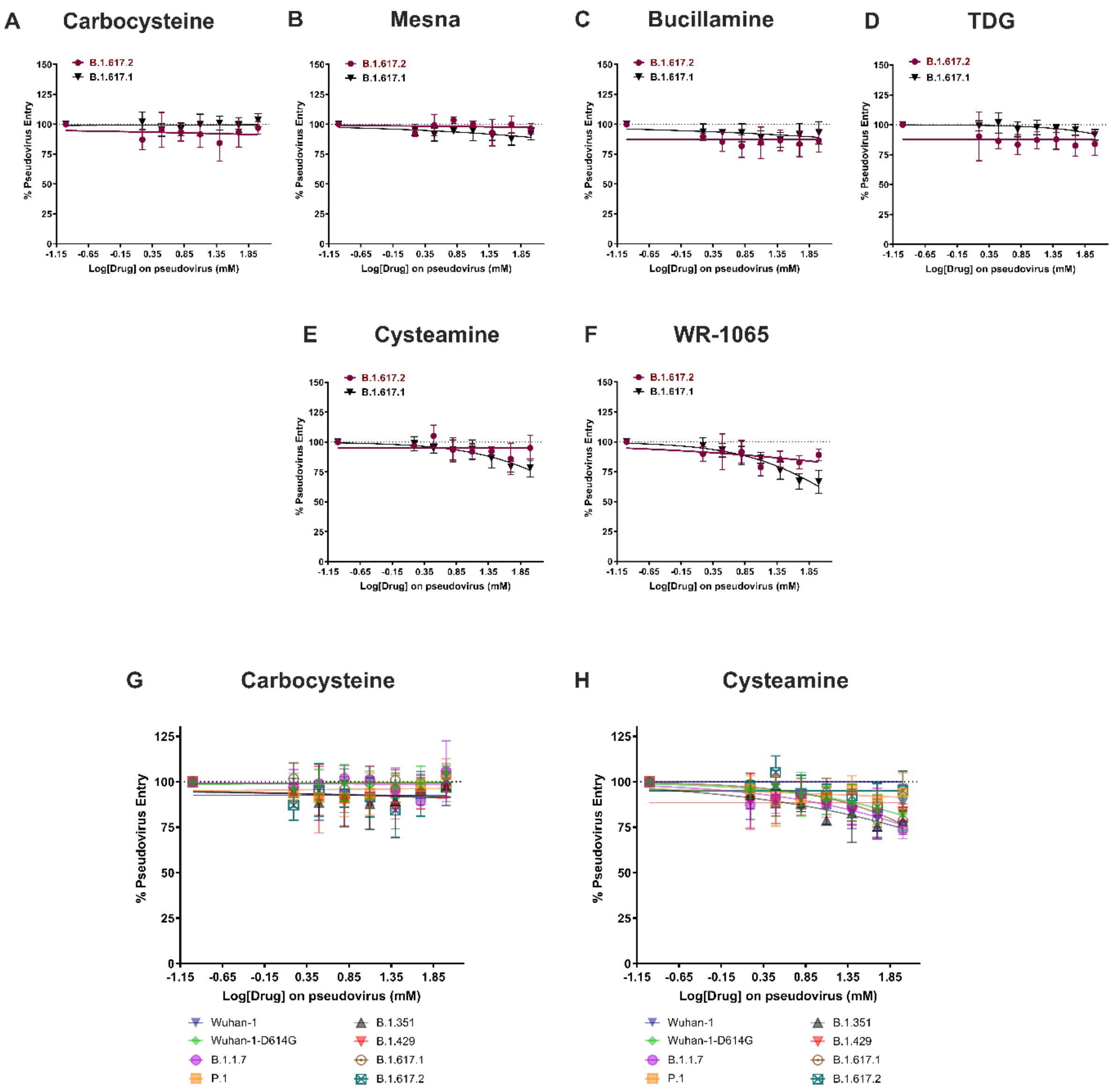


**Figure S5. Effect of pretreatment of 293T-ACE2-TMPRSS2 cells with thiol drugs on entry of variant pseudoviruses. (A to F)** Entry efficiency of B.1.617.1 and B.1.617.2 variant pseudoviruses, quantified by luciferase activity, when the cells were exposed to (A) carbocysteine (sulfide dug, negative control) and thiol drugs (B) Mesna, (C) bucillamine, (D) TDG, (E) cysteamine (F) WR-1065 prior to transduction with untreated virus (as illustrated in figure S2B), (n = 3-4). **(G, H)** Entry efficiency of the variant pseudoviruses, quantified by luciferase activity, when the cells were exposed to (G) carbocysteine and (H) cysteamine prior to transduction with untreated virus (n = 3-4). The X-axis are scaled to log10. Percentage changes are with respect to no drug control which is set as 100%. Data are mean ± SD.

**Figure S6.**

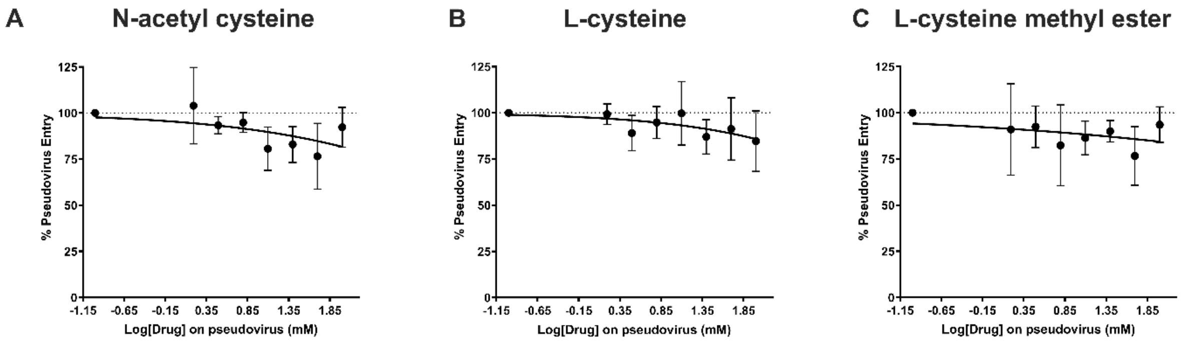


**Figure S6. Effect of pretreatment of 293T-ACE2-TMPRSS2 cells with cysteine derivatives on B.1.617.1 pseudovirus entry. (A to C)** Pseudovirus entry efficiency, quantified by luciferase activity, when the cells were exposed to cysteine derivatives (A) N-acetylcysteine, (B) L-cysteine, (C) L-cysteine methyl ester prior to transduction with untreated virus (n = 3-6). The X-axis are scaled to log10. Percentage changes are with respect to no drug control which is set as 100%. Data are mean ± SD.
